# Loop-extruder mediated rigidity can globally order bacterial chromosomes

**DOI:** 10.1101/2024.10.10.617531

**Authors:** Janni Harju, Till Armbruster, Chase Broedersz

## Abstract

Many bacterial chromosomes show large-scale linear order, so that a locus’s genomic position correlates with its position along the cell. In the model organism *E. coli*, for instance, the left and right arms of the circular chromosome lie in different cell halves. However, no mechanisms that anchor loci to the cell poles have been identified, and it remains unknown how this so-called “left-*ori*-right” organization arises. Here, we construct a biophysical model that explains how global chromosome order could be established via an active loop extrusion mechanism. Our model assumes that the motor protein complex MukBEF extrudes loops on most of the *E. coli* chromosome, but is excluded from the terminal region by the protein MatP, giving rise to a partially looped ring polymer structure. Using 3D simulations of loop extrusion on a chromosome, we find that our model can display stable left-*ori*-right chromosomal order in a parameter regime consistent with prior experiments. We explain this behavior by considering the effect of loop extrusion on the bending rigidity of the chromosome, and derive necessary conditions for left-*ori*-right order to emerge. Finally, we develop a phase diagram for the system, where order emerges when the loop size is large enough and the looped region is compacted enough. Our work provides a mechanistic explanation for how loop-extruders can establish linear chromosome order in *E. coli*, and how this order leads to accurate gene positioning within the cell, without locus anchoring.

## I. INTRODUCTION

Many bacterial species have a single circular chromosome, which is confined to a membraneless nucleoid within the cell. Physically, these chromosomes can be considered as highly compressed polymers, which often show a linear organization along the long axis of the cell: the genomic position of a locus correlates with its position along the nucleoid [1–5]. Disruption of this large-scale organization often has detrimental effects for chromosome segregation, which implies that maintaining chromosome order is of vital importance to bacteria [6–9].

In many bacterial species, the chromosome is organized so that the origin of replication (*ori*) is positioned at one cell pole, whereas the terminus of replication (*ter*), localizes at the opposite cell pole (Fig. 1(a)). This linear “*ori-ter* “ ordering can be maintained by anchoring of the origin of replication to a cell pole, and/or the Structural Maintenance of Chromosome (SMC) complex condensin [3]. Condensin is a loop-extruding motor protein that attaches onto DNA at a point and actively reels in a loop (Fig. 1(b)). In bacteria such as *Caulobacter crescentus* and *Bacillus subtilis*, condensins are loaded onto the chromosome near the *ori*, from where they proceed towards the *ter*, resulting in a queue of condensins “zipping up” the chromosomal arms [10, 11]. This effective linearization of the chromosome can be sufficient to maintain *ori*-*ter* organization, and can additionally enhance bacterial chromosome segregation during replication [12].

**FIG. 1.**
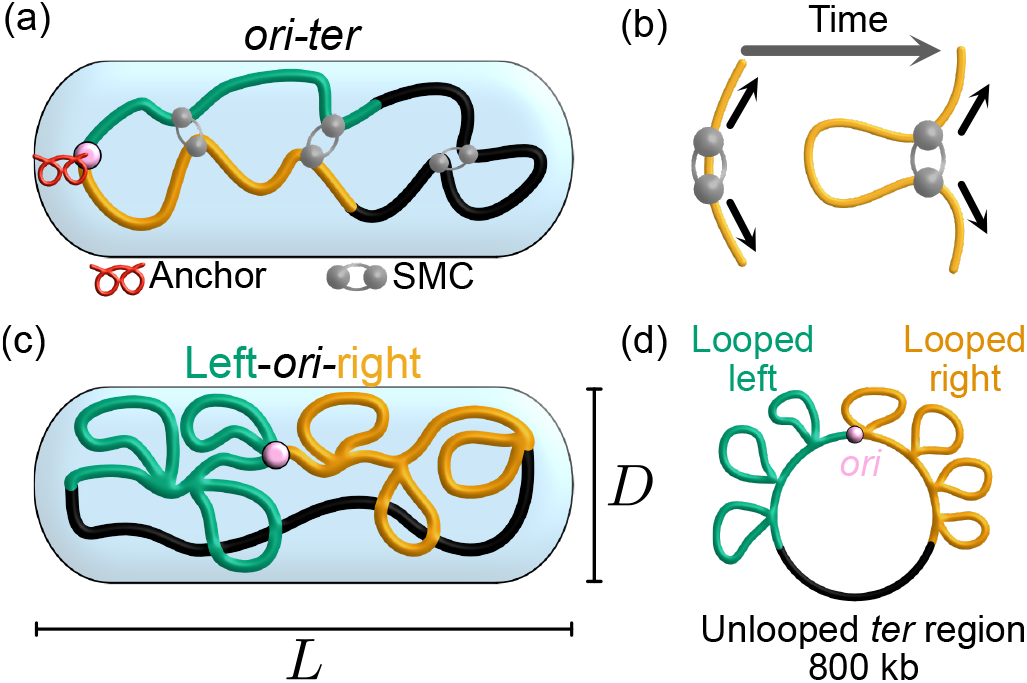
Bacterial chromosomes show linear order. **(a)** Coarse-grained cartoon of *ori*-*ter* ordered chromosome, as observed in species such as *Caulobacter crescentus*. The origin of replication is anchored to the cell wall, and loop-extruding condensins tie the arms of the chromosome together. **(b)** A loop-extruder attaches to DNA at a point. By moving both of its legs in opposite directions, it forms a loop. **(c)** Cartoon of a left-*ori*-right ordered chromosome, as observed in *E. coli*. We assume the chromosome is confined to a nucleoid of length *L* and diameter *D*. **(d)** Representation of the circular 4600 kb *E. coli* chromosome. Loop-extruders are excluded from the *ter* region by MatP.

Notably, the model organism *Escherichia coli* shows a different type of global chromosome order. In slowly growing *E. coli*, the origin of replication is located near the center of the nucleoid, and the left and right arms of the chromosome are located in opposite cell halves, in a “left-*ori*-right” configuration [13–16] (Fig. 1(c)). Experimental studies have shown that two protein complexes, MukBEF and MatP, are important for both the maintenance of left-*ori*-right order [17–19] and successful chromosome segregation in *E. coli* [20– Like bacterial condensin, MukBEF is an SMC complex that is thought to actively extrude loops. However, unlike bacterial condensin, MukBEF is not known to have a specific loading site. Instead, MukBEF’s distribution along the chromosome is controlled by MatP, which displaces MukBEF from a 800 kb region around the terminus of replication [23, 24]. In wild-type *E. coli*, we hence expect that 3800 kb of the chromosome is accessible to loop-extruding MukBEF complexes, whereas a 800 kb terminal region remains largely “unlooped” (Fig. 1(d)). However, a mechanistic explanation of how this partially looped structure gives rise to left-*ori*-right order is still lacking.

Motivated by imaging experiments that showed Muk-BEF clusters localizing mid-cell [17, 25–2 Murray & Sourjik proposed that MukBEF could organize into one or two clusters in the cell via a Turing patterning mechanism [28]. However, their model mainly focused on explaining nucleoid or *ori* positioning within the cell, rather than left-*ori*-right organization. More recently, Mäkelä & Sherratt performed 1D simulations of MukBEF loop extrusion in *E. coli* [24], but their work did not address how loop extrusion would affect the 3D organization of the chromosome. Finally, several coarse-grained polymer models that recapitulate *E. coli* chromosomal order have been constructed. In these models, left-*ori*- right organization is achieved by imposing constraints such as locus anchoring [29], static loops [30–32], attraction of loci within Macrodomains [30], or asymmetric coarse-graining of the chromosome in a concentric-shell model [33]. However, these models do not address what biological mechanisms could give rise to such constraints, and it therefore remains unclear how the global chromosome order emerges from molecular-scale processes.

In this work, we explain how unspecific loading combined with non-uniform off-loading of loop-extruders can establish left-*ori*-right order in an *E. coli* -like system. Using simulations of active loop-extrusion on a bead-spring polymer, we show that 50 MukBEF complexes with processivities of order 200 kb (consistent with experiments [18, 24, 34]) are sufficient to explain stable left-*ori*-right order as seen in experiments [19]. To theoretically explain how the density and processivity of loop-extruders control large-scale chromosome order, we propose that non-specifically loaded loop-extruders effectively rigidify a fraction of the chromosome, which then resists bending within the nucleoid. We predict a phase diagram for the system, where global chromosome order can emerge if the loops are large enough compared to the confinement width, and the looped region of the chromosome is sufficiently compacted compared to the confinement length. These predictions are confirmed using simulations. Finally, we find that our model can explain the accurate positioning of loci within the *E. coli* nucleoid [35]. Our work hence provides a mechanistic explanation for how MukBEF and MatP can maintain and establish left-*ori*-right order of the *E. coli* chromosome, illustrating how non-uniformly distributed loop-extruders can globally orient bacterial chromosomes.

## II. METHODS

### A. Computational model

We model the *E. coli* chromosome as a circular polymer with excluded volume interactions, consisting of 4600 coarse-grained monomers, each corresponding to 1 kb. The coarse-grained monomer size *b* is taken to be ≈ 30 nm, as used in previous studies [36, 37]. We assume that the chromosome is confined to a nucleoid of diameter *D* ≈ 1 µm and length *L* ≈ 1.8 µm [30, 38, 39] (Fig. 1(c)). We will also vary these dimensions to consider how cell growth would affect our model.

We assume that the chromosome consists of a terminal region of length *N*_*ter*_ = 800 kb and a non-terminal region of length *N* = 3800 kb (Fig. 1 (d)). A loop-extruder can bind to the chromosome at any point in the non-terminal region, and then reel in a loop by moving both of its legs in opposite directions (Fig. 1(b)). In the terminal region, we assume that loop-extruder binding is negligible, and that loop-extruders are unbound at an enhanced rate. Although the *E. coli* chromosome is known to consist of several macrodomains [40, 41], and highly transcribed genes can locally affect loop-extruder speeds [10, 11], for simplicity, we neglect inhomogeneities other than the enhanced off-loading of loop-extruders in the *ter* region.

To develop a computational model for our system, we adapt a previously used simulation scheme [42], where 1D simulations of loop extrusion are used to dynamically constrain 3D polymer simulations. The simulated loop-extruder trajectories are used to define the positions of moving harmonic springs that constrain loci in 3D bead-spring polymer simulations. The polymer is confined to a cylinder using harmonic potentials, and there is a finite excluded volume potential between beads. A more detailed description is provided in Appendix A.

In the 1D simulations, a fixed number *M* of loop-extruders can move on, bind to, or unbind from a periodic 1D lattice. The movement, binding, and unbinding rates are constant on the non-terminal part of the chromosome. In the *ter* region, the binding rate is set to zero, and the unbinding rate is increased by a factor of 100. Based on previous work in bacteria [37] and *in vitro* experiments with yeast condensin [43], we assume that upon head-on collision, two loop-extruder legs can bypass each other after a brief stalling period. Simulations without loop-extruder bypassing are also conducted for comparison (Appendix B).

We sample both the polymer and loop-extruder configurations at intervals corresponding to roughly 1-3 minutes (Appendix C), and calculate statistics once the system has relaxed (Supp. Fig. S1). This allows us to track chromosome orientation over time.

### B. Estimating MukBEF separation and processivity

Loop-extruder dynamics depend on two key parameters: the mean loop-extruder spacing, *d* = *N/M*, as well as the loop-extruder processivity, defined as λ = 2*v*_LE_τ, where *v*_LE_ is the mean loop-extruder leg speed and *τ* is the mean extrusion time. The amount by which loop-extruders can compact DNA has been shown to be determined by the ratio λ*/d* [44]. In the sparse regime (λ*/d* ≪ 1), loop-extruders rarely collide, and no significant compaction is given. When λ*/d* ≫ 1, the system reaches the dense regime, where most of the polymer is extruded into loops. Hence, the (relative) magnitudes of *d* and λ are expected to affect the behavior of our system, and we need to estimate their values based on experimental data.

The number of MukBEF complexes bound to DNA in cells has been approximated to be around 50 [18, 24], giving a loop-extruder separation of *d* = 3800*/*50 ≈ 75 kb. Estimates for the processivity of MukBEF, by contrast, have varied significantly. A Hi-C study of *E. coli* strains suggested that MukBEF enhances long-range contacts at the 280 kb scale [34]. On the other hand, Mäkelä & Sherratt reported a MukBEF association times of *τ* ≈ 60 s [24], giving processivities between 35-90 kb for loop extrusion speeds 0.6 kb/s [45] or 1.5 kb/s [10].

Given that these differing estimates for MukBEF’s processivity, we seek an alternative method of estimating λ. First, we note that for both λ = 30 and 280 kb, the dimensionless parameter λ*/d* is close to order unity, suggesting the system falls in the intermediate loop-extruder density regime. In this regime, the length of the polymer “backbone” not extruded into loops is approximately [44]

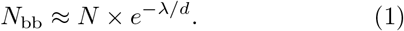

Upon over-expression of MukBEF, Mäkelä & Sherratt reported that an “axial core” of MukBEF, compacted by a factor *N/N*_bb_ = 1100 [24], become visible. Using that upon over-expression the loop-extruder number increased to *M* ≈ 176, we can use Eq. (1) to estimate λ ≈ *d ×* ln (*N/N*_bb_) ≈ 150 kb. Given these considerations, we will run our simulations for λ between 30 and 300 kb.

## III. RESULTS

### A. Loop-extruders can maintain and establish left-*ori*-right order

To determine whether loop-extruders could explain the stable left-*ori*-right order observed in WT *E. coli*, we start by comparing simulations with no loop-extruders, with no unlooped *ter* region, and with both loop-extruders and an unlooped *ter* region (referred to as the “WT model”). We initialize all simulations from a left-*ori*-right configuration, and then track whether this orientation is maintained.

Animations and snapshots reveal that in simulations without loop-extruders or without an unlooped *ter* region, the left and the right arm of the chromosome mix (Fig. 2(b), Supp. Video 1,2). By contrast, WT simulations with *M* = 50 and λ = 200 appear to maintain left-*ori*-right order, with the arms of the chromosome staying in the nucleoid half that they started in (Supp. Video 3).

**FIG. 2.**
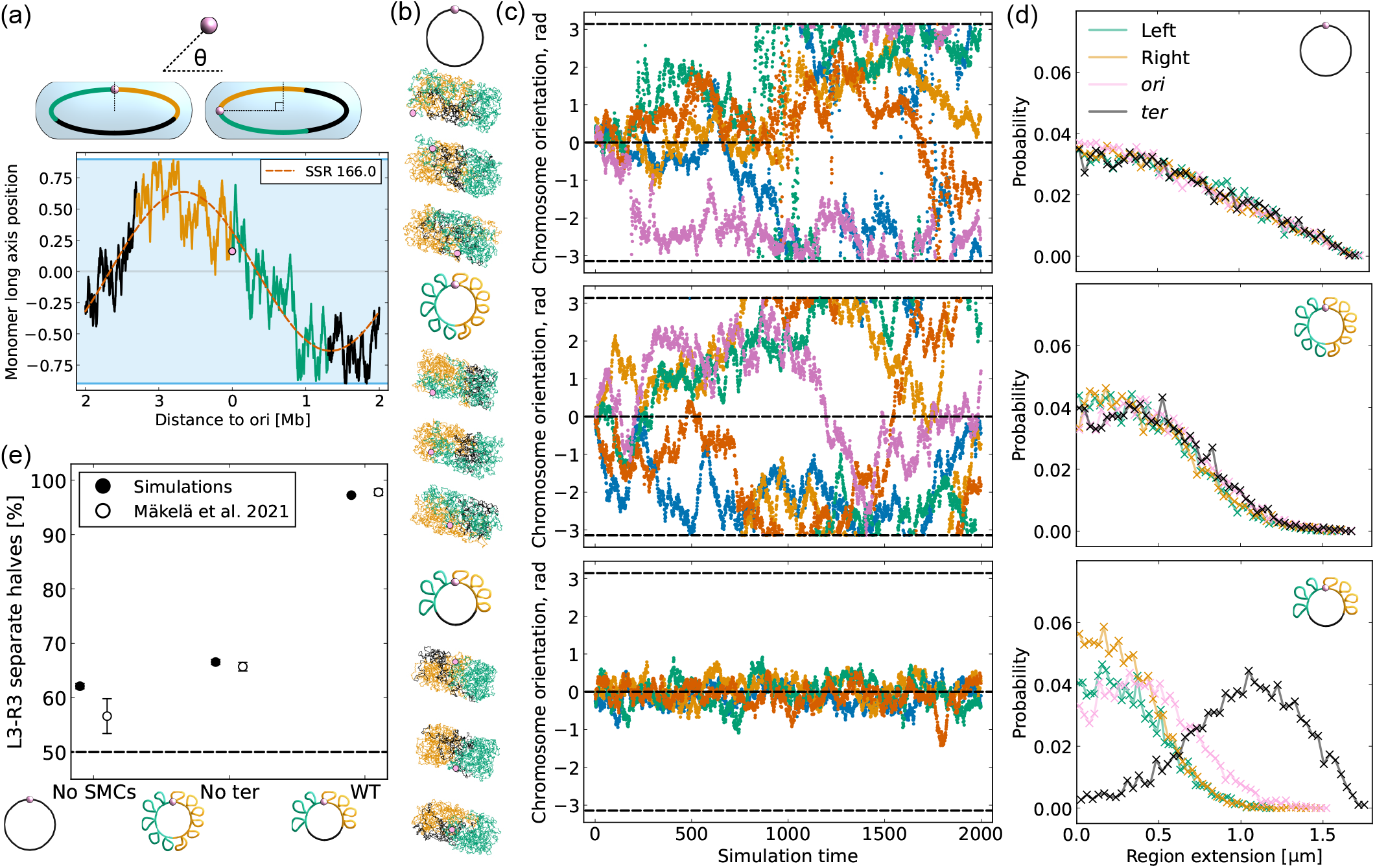
Simulations show stable left-*ori*-right order. **(a)** The orientation angle for a chromosome configuration is found by fitting the monomer long axis position curve with a sinusoidal function. The phase shift from left-*ori*-right organization defines *θ*. Blue band in figure indicates confinement height. Sum of squared residuals (SSR) is shown. Supp. Fig. S2 shows more example curves. **(b)** Snapshots from simulations without loop-extruders, without an unlooped *ter* region, as well as WT simulations. Icons describe models. In snapshots, the looped left and right arm of the chromosome are in green and orange, whereas the 800 kb terminal region is in black. **(c)** Orientation angles over time for the same models. Five simulation trajectories are shown. Dashed lines indicate 0, *±π* rad. **(d)** Distributions of long-axis extensions of 800 kb regions for the same models. The regions are centered at the origin, the terminus, and the mid-points of the left and right arm. **(e)** Comparison of probability of loci L3 and R3 being in the same half of the nucleoid, compared to experimental data from [19].

To parametrize the orientation of the chromosome within the confinement, we define an “orientation angle” *θ*. Briefly, left-*ori*-right configurations are assigned an orientation angle of *θ* = 0 rad, *ori*-*ter* configurations *θ* = *π/*2 rad, and so forth (Fig. 2(a), Appendix D). The orientation angle provides a convenient characterization of the organization of a single chromosome configuration. We find that in simulations without loop-extruders or without an unlooped *ter* region, the orientation angle diffuses across all orientations (Fig. 2(c)). In both cases, the extensions of 800 kb regions centered at 0^°^, 90^°^, 180^°^and 270^°^ relative to the *ori* also have identical distributions (Fig. 2(d)), further illustrating that these models show rotational symmetry. Furthermore, these results confirm that the duration of our simulations (equivalent to roughly 33-100 h, Appendix C) is long enough to show significant changes in chromosome orientation.

In the WT model with *M* = 50 and λ = 200 kb, all simulation trajectories retain a left-*ori*-right organization for the duration of the simulations (Fig. 2(c)). However, we find that this order is only maintained for high enough processivities; for λ = 25 − 50, simulations show significant rotation, whereas for λ = 100−150, we observe rapid “flips” from a left-*ori*-right to a right-*ori*-left orientation (Supp. Fig. S3). *In vivo*, the relative long axis positions of L3 and R3 loci (at −128^°^ and 122^°^) have been observed to flip every 12.5 cell cycles, approximately 10 times less frequently than in MatP lacking cells [19]. Therefore, occasional flipping behavior in our simulations could still be consistent with experimental observations. In our simulations, with *M* = 50, λ = 150 kb, the no *ter* region model has a 7.5 times higher L3-R3 flipping frequency than the WT model, whereas for *M* = 50, λ = 200, the flipping frequency is 20 times higher. We hence conclude that in our simulations, processivities between λ = 150 − 200 kb give similar levels of left-*ori*-right order stability as observed in experiments.

By calculating the fraction of configurations where the L3 and R3 loci are in different nucleoid halves, we can further compare our simulations to experimental data from [19]. We find that in WT simulations, the two loci are in different cell halves over 95% of the time, in excellent agreement with experiments (Fig. 2(e)). This suggests that our simple simulation model gives rise to similar MukBEF and MatP dependent chromosome organization as seen in experiments, at least for high enough processivities λ ≈ 200 kb.

Finally, we run WT simulations starting from an *ori-ter* configuration to test whether loop-extruders can not only maintain, but also establish left-*ori*-right organization. We find that left-*ori*-right order is established in approximately 100 simulation steps (roughly 100-300 min, Appendix C), comparable to the time it takes chromosomes to rotate by 90^°^in simulations without an unlooped *ter* region (Fig. 2(b)). This newly established order is then maintained for the rest of the simulation (Supp. Fig. S4).

We conclude that based on our simulations, 50 bidirectional loop-extruders can give rise to stable left-*ori*-right order in an *E. coli* -like system, using reasonable values for λ and the cell size.

### B. Theoretical conditions for stable left-*ori*-right order

Next, we develop a theoretical framework to explain how loop-extruders can maintain and establish left-*ori*- right order. When enough loop-extruders are present, the looped region of the chromosome resembles a polymer brush, consisting of loops emanating from a polymer backbone (Fig. 3(a)). Since excluded volume interactions between side-chains can give rise to an entropic cost to bending [46, 47], we expect that the looped region will have a higher bending rigidity than the unlooped *ter* region. This partially rigid ring structure can favor left-*ori*-right configurations where the looped region is extended along the long axis of the cell. However, there are three necessary conditions for such a configuration to be favored: first, the unlooped *ter* region’s contour length must be long enough to stretch across the nucleoid; second, the loops must be large enough to sufficiently rigidify the chromosome; and third, the looped region has to have enough space to extend within the confinement. We will now express these conditions more formally.

**FIG. 3.**
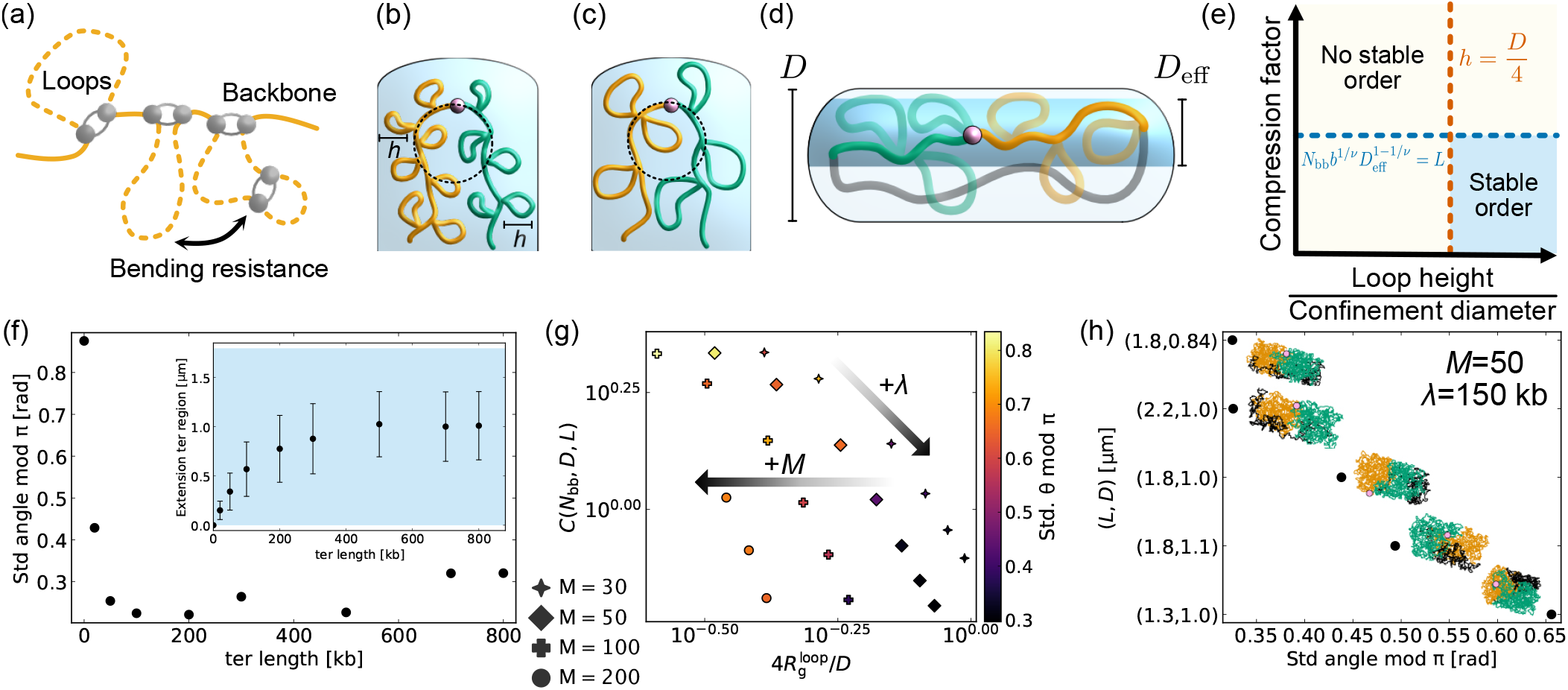
Conditions for left-*ori*-right stability. **(a)** The looped region consists of loops (dashed lines) and a polymer backbone (full line). **(b)** The looped arms of the chromosome can lie parallel to each other without loop deformation if the loop height *h* is smaller than *D/*4. **(c)** If *h > D/*4, bending the looped region causes loop deformations. **(d)** We assume that the presence of loops and the unlooped region reduces the effective diameter of confinement for the backbone. **(e)** A sketch of a phase diagram for the system. Stable left-*ori*-right order is expected to emerge if the loops are large enough compared to the confinement diameter, and that the backbone is sufficiently compacted compared to the confinement length. **(f)** The standard deviation of the orientation angle modulo *π* (Appendix D) for different *ter* lengths. Inset: The extension of the *ter* region as a function of *ter* region length. Blue color indicates confinement length. **(g)** A scatter plot of the standard deviation of the orientation angle modulo *π* for simulations with *M* = 30, 50, 100, 200 loop-extruders, with λ between 25 kb and 300 kb. Arrows indicate directions of changing λ with constant *M* and changing loop-extruder number *M* with fixed λ*/d*. Plots for other stability measures are shown in Supp. Fig. S6. **(h)** Comparison of *θ* mod *π* for simulations with *M* = 50, λ = 150 kb, but varying dimensions. Longer and narrower confinements improve left-*ori*-right organization. Insets show simulation snapshots.

First, we note that a left-*ori*-right configuration is only possible if the unlooped terminal region can stretch across the nucleoid. A terminal region of *N*_*ter*_ = 800 with *b* = 0.03 µm has a contour length of 24 µm, much larger than the cell length. We hence expect that in WT *E. coli*, the length of the terminal region is more than sufficient for left-*ori*-right order to be established.

Second, we consider how large loops would need to be to sufficiently rigidify the chromosome for left-*ori*-right organization to become favorable. Intuitively, a polymer brush has an increased bending rigidity compared to an unbranched polymer, because upon bending, the side-chains of the brush are pushed together. Due to excluded volume interactions, this introduces an entropic cost to bending. This intuitive picture suggests that the effective side-chain height *h* of the brush sets the length scale at which side-chain interactions start to disfavor bending. Scaling analyses confirm that the persistence length *ℓ*_p_ of a bottle-brush polymer scales linearly with the brush height *h* [48], or as *ℓ*_p_ ∼ *h*^2^ for much longer side chains [46, 49].

Consider an evenly looped polymer with loops of height *h*, confined to a tube of diameter *D*. For the polymer to bend around within the confinement tube, the radius of curvature must be less than or equal to *D/*2−*h* (Fig. 3(b-c)). These bends become thermodynamically inaccessible when the effective persistence length of the brush is of the same order of magnitude: *ℓ*_p_ ∼ *h* ∼ *D/*2−*h*, or *h* ∼ *D/*4. We also note that when *h* ≤ *D/*4, two looped arms of the chromosome can lie side by side within the confinement without significant loop overlap. We hence expect *h/D* ≈ 1*/*4 to define an approximate minimum loop height at which left-*ori*-right organization can become stable.

Finally, even if the loops on the chromosome are sufficiently large to inhibit bending inside the confinement tube, if the confinement length *L* is too short, the looped region may be forced to buckle. If the looped region has rigidified sufficiently to behave as a semi-flexible polymer, we expect buckling to occur at strains of order tens percent [50]. Therefore, for stable left-*ori*-right order to emerge, the looped region’s rest length in the tube should be shorter than or comparable to the confinement length.

As a first approximation for the rest length of the looped region, we neglect the effects of the loops, and only consider a backbone of *N*_bb_ monomers. Following De Gennes [51], the expected extension of a linear polymer in a tube, with excluded volume interactions, can be calculated using a simple blob argument, upto a prefactor of order unity. Below length scales of order *D*, a polymer segment with *n* monomers is effectively unconfined: the radius of gyration of the segment scales as *R*_g_ *∝ n*^*ν*^*b*, where *ν* is the Flory exponent, *ν* ≈ 3*/*5 for real polymers in 3D. This means that the polymer effectively splits into “confinement blobs”, inside of which the polymer is effectively unconfined. The number of monomers per blob, *g*, can be found by setting the blob radius of gyration equal to the confinement diameter: *g* ∼ (*D/b*)^1*/ν*^. Due to excluded volume interactions, the blobs align sequentially along the confinement tube, so that the extension *z* of the polymer is given by the number of blobs times the blob size *D*:

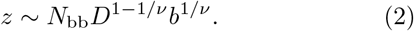

Based on this calculation, we might hence expect the backbone to be sufficiently compacted when

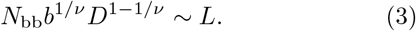

The simple argument above neglects the fact that the loops and unlooped terminal region reduce the accessible volume for the backbone, effectively forcing it to straighten. To correct for this effect, we start by considering a left-*ori*-right configuration, where the backbone, loops and terminal region all stretch out across the same length. We then seek to define an effective confinement diameter *D*_eff_ for the backbone [52] (Fig. 3(d)). We assume that the fraction of the volume occupied by the backbone scales with the fraction of monomers in the backbone: 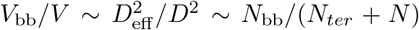.The effective diameter of the backbone’s confinement is then given by

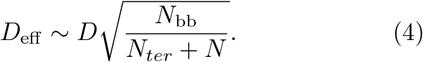

Replacing *D* by *D*_eff_ in Equation (3), and dividing by *L*, we find a condition

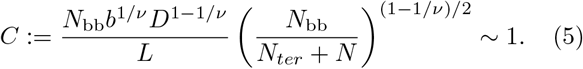

We name this dimensionless parameter the compression factor *C*, since it describes the fraction by which the backbone (and looped region) would have to be compressed in order to fit into a tube of length *L* and diameter *D*_eff_. Buckling is expected to occur when *C* ≫ 1.

Eq. (5) features two contributions of the backbone length *N*_bb_. The first linear term simply describes how a longer backbone needs a longer confinement length to stretch out. The second contribution describes a trade-off between compaction and accessible volume; at higher levels of backbone compaction, more of the chromosome is extruded into loops, which reduces the effective diameter of the tube the backbone can extend in. We confirm our scaling prediction 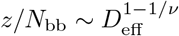 using simulations of a chromosome in an infinitely long confinement tube (Appendix E, Supp. Fig. S5).

Our theoretical arguments hence suggest that we expect the stability of left-*ori*-right order in *E. coli* to be controlled by two dimensionless parameters; the size of the loops compared to the confinement diameter (*h/D*), as well as the expected extension of the backbone compared to the length of the confinement (*C*, given by Eq. (5)). We hence propose that the system can be described by a phase diagram of (*h/D, C*) (Fig. 3(e)). In the left-side of the diagram, with *h/D* ≪ 1, the loops are too short to prevent bending within the confinement tube. In the top-half of the diagram (*C* ≫ 1), the looped region is too long to extend across the nucleoid. Hence, we only expect stable left-*ori*-right order to emerge in the bottom-right part of the diagram, where loops are large enough to rigidify the chromosome, and the looped region is compacted enough to extend across the nucleoid. If these two conditions are met, a partially looped chromosome structure can give rise to stable left-*ori*-right chromosome organization.

### C. Estimating control parameters *h* and *C* in *E. coli*

Based on the values *M* ≈ 50 and λ ≈ 30 − 300 kb, we now try to approximate the loop height and backbone extension in the relevant parameter range.

In the intermediate loop-extruder density regime (λ*/d* ∼ 1), the average genomic length of a loop is of order *ℓ* ≈ λ (Supp. Fig. S5), although stalling upon loop-extruder collisions somewhat decreases *ℓ* (Appendix B). To relate this mean loop length to the expected height *h* of the loops, we must consider whether the system falls into a dense bottlebrush regime, where loops are tightly packed and extend radially from the backbone, or into the “mushroom regime”, where the spacing between loops is comparable to their radius of gyration [49]. In the mushroom regime, we expect loops to show their usual radius of gyration scaling, so that *h* ∼ λ^*ν*^*b*. By contrast, in the dense bottlebrush regime, we expect a stronger scaling of *h* with λ, as well as a dependence on the loop spacing [53, 54].

Our simulations suggest that 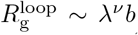,with *ν* ≈ 0.4, largely independent of the loop spacing (Supp. Fig. S5). This suggests that the simulations fall into the mushroom regime, as can also be argued based on the parameter range (Appendix F). The found exponent *ν* is consistent with previous reports of *ν* ≈ 0.39 − 0.45 for solutions of ring polymers [55–59], although at higher ring densities *ν* ≈ 1*/*3 is expected [60, 61].

Using 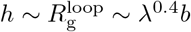,for λ = 150 − 300 kb, we find loop heights of order *h* ∼ 200 − 300 nm. Since for a cell of diameter 1 µm, we expect stability to require loops of size *h* ≈ 250 nm, these processivities could hence lead to sufficiently large loops for stable left-*ori*-right order.

Finally, we ask whether loop-extruders can give rise to sufficient compaction for stable left-*ori*-right order. Using Eq. (1), for processivities of λ = 30, 150 or 300 kb, we would find backbone lengths *N*_bb_ ≈ 2500, 530, 70 kb correspondingly. For *D* = 1 µm and *L* = 1.8 µm, this would give compression factors of *C* ≈ 5.7, 2.1 or 0.5. Hence, at least for larger processivities, loop extrusion could compact the looped region sufficiently for it to extend across the nucleoid.

In summary, we find that a mean separation of 75 kb, as well as a processivity of 150 kb or more, should yield both sufficiently large loops and sufficient backbone compaction for left-*ori*-right order to emerge. This is consistent with our initial simulation results (Section III A), which showed stable chromosome orientations for *M* = 50 and λ = 200 kb (Fig. 2), and occasional flipping for *M* = 50 and λ = 150 kb.

### D. Unlooped *ter* region of 50 kb can be sufficient for left-*ori*-right order

We now wish to use our simulation model (Section II A) to test our predicted criteria for the emergence of chromosome order. We start by testing whether shortening the unlooped *ter* region can destabilise left-*ori*- right organization. We hence keep *N* = 3800, λ = 200 kb, *M* = 50 and the confinement dimensions fixed, but reduce the length of the unlooped *ter* region, *N*_*ter*_, in our simulations. For terminal regions of *N*_*ter*_ = 500 kb or more, the looped region has similar levels of mean extension, whereas for *N*_*ter*_ *<* 500 kb it starts to contract (Inset, Fig. 3(f)). Using Eq. (2), this transition can be understood as the point when the expected extension of the terminal region becomes comparable to the cell length: 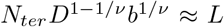,suggesting a critical value of *N*_*ter*_ ≈ 550 kb.

Notably, terminal regions much shorter than 500 kb still allow the system to maintain a stable chromosome orientation, all the way down to *N*_*ter*_ = 50 kb (Fig. 3(f)). At this point, the contour length of the unlooped *ter* region is barely sufficient for the region to stretch out across the nucleoid (50 *×* 0.03 = 1.5μm, compared to a nucleoid length *L* = 1.8μm), but despite the entropic cost of this extension, left-*ori*-right order is maintained. For *N*_*ter*_ = 20 kb, organizational stability decreases. We hence conclude that an unlooped *ter* region much shorter than 800 kb would also be sufficient to maintain stable chromosome order, provided that the region’s contour length is at least as long as the nucleoid.

### E. Both compaction and large enough loop-size required for stability

Next, we test our predictions for how the loop size and the backbone compaction affect left-*ori*-right order. We hence conduct simulations where we vary the number of loop-extruders *M* (or equivalently *d*) and the processivity λ. We then use the mean loop radius of gyration, 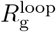,as a proxy for the loop height *h*, and calculate the compression factor *C* using the the confinement dimensions and the mean value of *N*_bb_.

Plots of stability measures as a function of 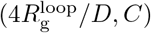(Fig. 3(g), Supp. Fig. S6) resemble our predicted phase diagram (Fig. 3(e)): stable chromosome organization requires both 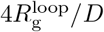 and *C* close to unity. Furthermore, plots of stability measures as a function of only the loop-size, only the backbone length, or only the loop height do not show a clear trend (Supp. Fig. S7), suggesting that both backbone compaction and a sufficient loop size are necessary for stable left-*ori*-right order.

For approximately the same *N*_bb_ (horizontal line in Fig. 3(g)), smaller *M* is predictive of stabler organization, as expected since the loop size increases. For fixed *M*, increasing the processivity λ increases the loop size and thereby left-*ori*-right stability (diagonal line in Fig. 3(g)). However, strikingly, for simulations with *M* = 200, although the backbone of the chromosome is compacted, the loop-sizes remain insufficient to give rise to stable left-*ori*-right order in our simulations (circles in Fig. 3(g)). We note that a condition on the loop height (*h* ∼ *D/*4), together with the fact that at high loop-extruder densities the mean loop-size decreases [44], gives rise to a maximum number of loop-extruders that can maintain left-*ori*-right order. A simple scaling argument suggests that this maximum loop-extruder number is of order 190 (Appendix G), consistent with our simulations.

Next, we keep λ and *d* fixed, and adjust the confinement dimensions. For increasing *L*, we expect *C* ∼ *L*^−1^ to decrease, and left-*ori*-right order to stabilize. For increasing *D*, on the other hand, although *C* ∼ *D*^1−1*/ν*^ still decreases, *h/D* now also decreases, which should destabilize chromosome organization. Consistent with these predictions, we find that increasing the confinement length in our simulations can stabilize chromosomal order, whereas increasing the confinement diameter leads to destabilization (Fig. 3(h), Fig. S8). The results from these simulations can be placed in the same phase diagram as before, showing that 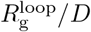 and *C* are still predictive of left-*ori*-right stability (Supp. Fig. S6, S8), even though the density of the system changes.

In summary, simulations confirm our theoretical predictions that both large enough loops and sufficient backbone compaction are required for the maintenance of left- *ori*-right order via loop extrusion. We also find that longer and/or narrower cells should show more stable left-*ori*-right organization.

### F. Left-*ori*-right order leads to accurate positioning of loci

We have hence demonstrated that loop-extruders can stably organize the *E. coli* chromosome, but what functions could this organization serve? In addition to potentially enhancing chromosome segregation [6, 8], we note that orienting the circular chromosome leads to accurate positioning of genes within the nucleoid. In *E. coli*, ribosomes segregate from the nucleoid to the cell poles [62, 63], and it has been suggested that chromosome organization could be linked to transcription or translation [64, 65]. Additionally, the information stored in gene positions could be used to guide intra-cellular processes [66].

Experiments have shown that the long axis positions of most *E. coli* loci follow approximately Gaussian distributions, with standard deviations ≈ 10% of the cell length [35]. Loci in a ∼ 370 kb “crossing region” around the terminus, on the other hand, were found to either localize at either nucleoid edge, or then uniformly across the length of the nucleoid. We now ask whether our model could explain these results.

We find that in our WT model, the standard deviation of locus positions in the looped region is approximately 10-20 % of the nucleoid length (Fig. 4(a)). In the unlooped *ter* region, by contrast, the distributions show standard deviations of approximately 20-30% the nucleoid length, similar to levels seen in simulations without loop-extruders or without an unlooped *ter* region. We find that across our simulations, more stable chromosome organization is correlated with more accurate locus positioning (Fig. 4(b)). This shows that left-*ori*- right order can lead to accurate gene localization within the looped region.

**FIG. 4.**
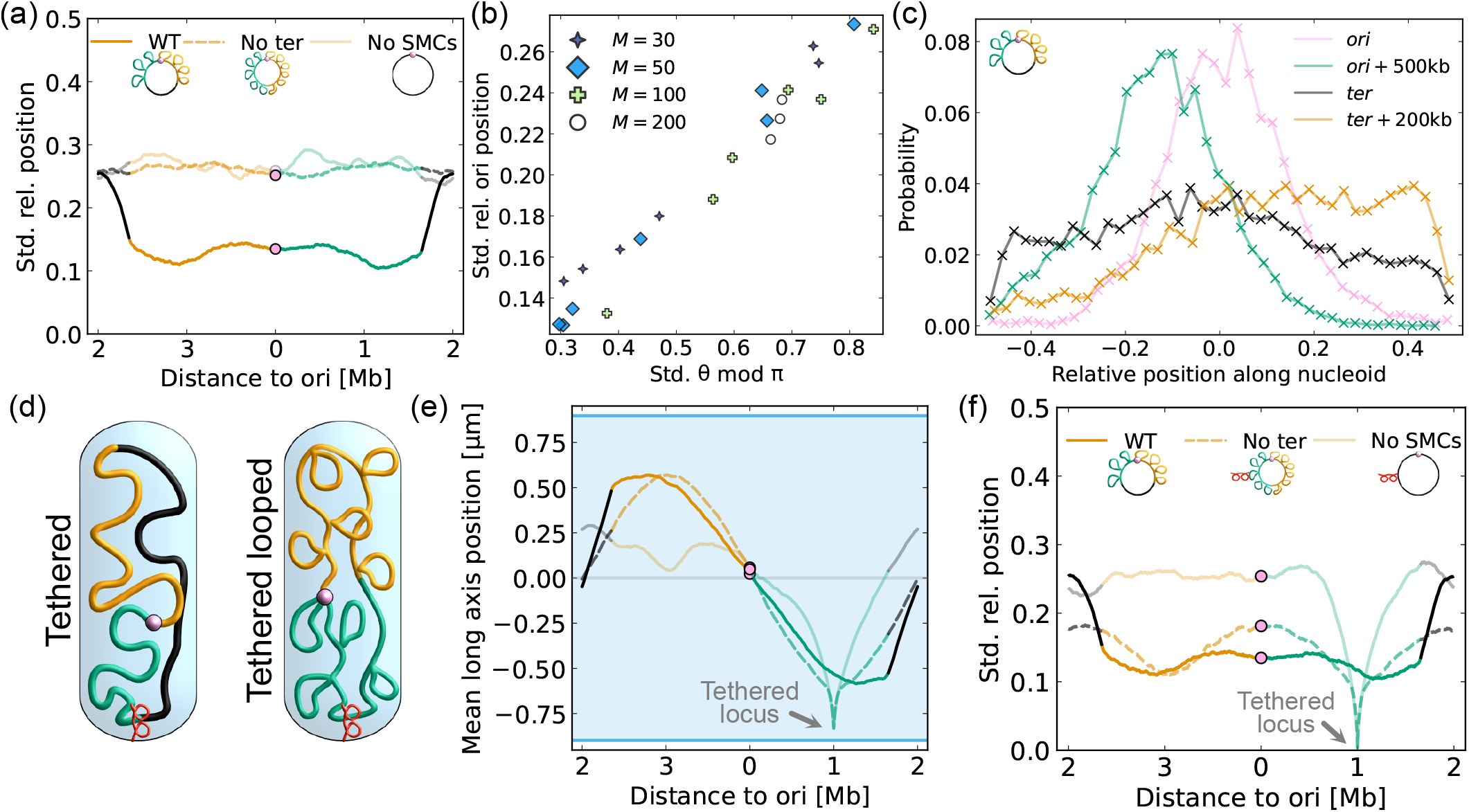
Left-*ori*-right order allows accurate positioning of loci. **(a)** Standard deviation of the long axis position relative to confinement length as a function of genomic position, for WT, no unlooped *ter* region, and no SMCs simulations. Same parameters as shown in Fig. 2. **(b)** The standard deviation of the *ori* ’s long axis position relative to the confinement length as a function of left-*ori*-right order stability, from simulations with fixed *D* and *L* but varying *M* and λ. **(c)** Histogram data for the positioning of various loci in simulations with *M* = 50 and λ = 200 kb. **(d)** In simulations with anchoring, the locus at 90^°^to the *ori* is tethered to the nucleoid pole with a harmonic potential. We consider simulations without loop-extruders or without an unlooped *ter* region (*M* = 50, λ = 200). **(e)** Mean long axis positions of loci as a function of genomic position. Blue background indicates confinement length. Legend in subfigure (f). Without loop-extruders (faint line), the untethered chromosomal arm (orange) does not show linear order. **(f)** Similar to subfigure (a), but for simulations with the monomer at 90^°^(indicated by arrow) tethered to the nucleoid pole. WT simulations were conducted without a tether.

To further compare our simulations to data from [35], we plot histograms of the long axis distributions of several loci (Fig. 4(c)). In simulations with stable left-*ori*-right order, these histograms are consistent with the experimental data; loci around the origin have approximately normal long axis distributions, whereas loci close to the *ter* show a relatively uniform distribution. These results suggest that our model captures both the accurate positioning of loci near the origin, as well as the more random positioning of loci in the terminal region.

Some bacterial species achieve chromosome order by anchoring a specific locus to a cell pole. To test whether loop-extruder mediated chromosome order leads to more or less accurate gene positioning than anchoring, we run simulations with the locus at 90^°^ to the origin tethered to the edge of the nucleoid (Fig. 4(d)). We find that, without loop-extruders, the chromosome is uncompacted, and tethering a single locus to the cell pole is insufficient to maintain linear order of the chromosome (Fig. 4(e), Supp. Fig. S9), consistent with previous reports [29]. We therefore tether the same locus in simulations with loop-extruders but no unlooped *ter* region. In these simulations, the chromosome is uniformly compacted, and tethering a single locus introduces linear order of the chromosome (Fig. 4(e)). However, the standard deviation in the relative position of the *ori*-proximal regions is higher than in WT simulations without anchoring (Fig. 4(f)). This indicates that loop-extruder mediated left-*ori*-right order can give rise to more accurate gene positioning than anchoring a single locus to the cell pole.

## IV. DISCUSSION

In conclusion, using biophysical theory and extensive polymer simulations, we have shown that non-uniformly distributed loop-extruders can maintain and establish left-*ori*-right order in unreplicating *E. coli*. Our work hence provides a mechanistic explanation for experiments which have shown that both MukBEF or MatP are necessary for chromosomal order in *E. coli* [15, 18, 19, 67]. Our work suggests that the formation of sufficiently large loops (∼ 200 kb) is critical for establishment of stable left-*ori*-right order. Since the distribution of loop sizes formed by passive mechanisms decays quickly with a power-law dependence [68, 69], an active loop formation process seems necessary for the creation of loops of this size. Although MukBEF processivities of order 200 kb are consistent with Hi-C data [34], as well as the level of chromosome compaction (Section II B), they appear inconsistent with reported MukBEF association times of order a minute [24, 70]. Further studies are hence needed to confirm whether and how MukBEF could form large enough loops for stable chromosome organization. We also note that although our work has focused on an ideal case where loops are solely formed via loop extrusion, the organization mechanism we propose could be enhanced by other looping mechanisms. The main role of active loop extrusion in our model is to give rise to a sufficient coverage and correct distribution of large enough loops on the chromosome; simulations with fixed loop-extruder positions also show stable left-*ori*-right organization (Supp. Fig. S10, Appendix H).

The length of the terminal region, unlike the loop size, is not critical to the maintenance of left-*ori*-right order in our model; even a 50 kb unlooped region could be sufficient to stretch out across the nucleoid. One might hence wonder why the *E. coli* terminal region is as long as 800 kb. We note that stretching a terminal region beyond its rest length comes with an entropic cost. Furthermore, if the size of the genome is fixed, a smaller number of loop-extruders is needed to compact a shorter looped region; a longer unlooped terminal region could hence require less resources for maintaining left-*ori*-right order.

Future work could explore how loop extrusion during replication affects *E. coli* chromosome segregation. The interplay between loop extrusion and replication in *E. coli* is complicated by several factors, such as multifork replication [71], and potential interactions between Muk-BEF and topoisomerase IV [72–74]. Nevertheless, since previous simulation work has suggested that deterministically placed fixed loops can enhance bacterial chromosome segregation [31], and since a looped polymer structure due to SMCs condensin I and II facilitates mitotic chromosome segregation in eukaryotes [47], loop extrusion by MukBEF is a plausible segregation mechanism in *E. coli*.

Our work suggests multiple experimentally testable hypotheses. First, we predict that altering the nucleoid size [75] can impact the stability of chromosomal order; wider nucleoids should show less stable left-*ori*-right organization, whereas longer nucleoids should show more stable organization. Second, the orientation of the chromosome is determined by the position of the unlooped *ter* region. Therefore, mutants with relocated MatP binding sites [67], should show a rotated chromosome organization. Third, perhaps counter-intuitively, if the number of loop-extruders is increased to *M* ∼ 200 [24], we expect left-*ori*-right order to be destablized.

Finally, our model shows that loop-extruder mediated left-*ori*-right ordering of chromosomes leads to accurate localization of genes along the nucleoid’s long axis, without a need for locus anchoring. Although the possible functional benefits of such linear chromosome ordering remain unclear, this illustrates that bacteria have evolved diverse mechanisms for locus positioning within the nucleoid.

## ACKNOWLEDGMENTS

We thank Joost de Graaf for discussions. We would also like to thank Jarno Mäkelä for discussions, suggestions, and feedback. This project has received funding from the European Research Council (ERC) under the European Union’s Horizon 2020 research and innovation programme (Grant agreement No. 101122863).

## DATA AVAILABILITY

All code used for simulations and data analysis can be found at github.com/PLSysGitHub/e coli loop extrusion.

All simulated data can be found at Zenodo.

## Appendix A: Simulations

For 1D simulations of loop extrusion, we adapt the looplib package [42] as follows: a periodic lattice with *N*_total_ = 4600 is used to simulate the circular bacterial chromosome; and bypassing of loop-extruders upon collision is implemented, since this is consistent with previous modeling works in bacteria [37] as well as *in vitro* observations of yeast condensins traversing each other [43]. Briefly, when two loop-extruder legs collide head-on, ie. travelling in opposite directions, they are allowed to traverse each other at a reduced rate.

The number of active loop-extruders during a simulation is kept fixed. To model the *E. coli* chromosome, we assume a constant loading probability of loop-extruders everywhere except the terminal region, where the loading probability is zero. The off-loading rate is set by the extrusion time *τ* of the loop-extruders, proportional to the processivity λ. In the terminus region, the off-loading rate is increased by a factor of 100. This models an ideal case where there is a negligible amount of loop-extruders present on the terminal region.

For 3D polymer simulations, we use the polychrom wrapper [76] for the molecular dynamics library OpenMM [77]. The bead-spring polymer is confined to a cylinder using a harmonic potential. Excluded volume interactions between beads are modeled by a repulsive potential

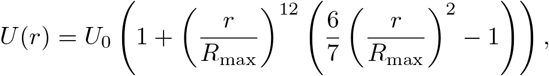

where *U*_0_ = 5 k_B_T is the depth of the potential; *r* is the separation between two beads; and *R*_max_ is the distance at which the potential’s derivative becomes zero (1.05 times the spring length *b* = 30 nm). This potential allows occasional strand passing, mimicking the effect of topoisomerases *in vivo*, and our simulations hence do not conserve the chain topology. However, the potential is more than sufficient for linear chains to swell and show the appropriate Flory exponent of *ν* = 3*/*5 [12].

A 3D simulation time-step consist of updating the positions of additional springs representing loop-extruders based on the 1D simulations, and then allowing the polymer to relax.

For each set of simulation parameters, at least five simulations are conducted. Unless otherwise specified, simulations are started from a left-*ori*-right orientation.

Based on the convergence of the mean long axis positions of loci as well as the region extensions, we conclude that most simulations converge after 250-500 simulation time-steps (Supp. Fig. S1), equivalent to roughly 1-8 hours (Appendix C). For all averaged data presented, the mean is calculated over all time-points after 1000 time-steps.

## Appendix B Loop-extruder traversal upon collision

*In vitro* studies have shown that, upon collision, yeast condensins can traverse each other after a stalling time *τ*_stall_ of approximately 7 seconds [43]. In addition, a simulation study focused on the bacterium *B. subtilis* found that patterns on Hi-C maps of mutant strains could only be explained if loop-extruding condensins were able to traverse each other after *τ*_stall_ ≈ 20 *±* 10 s [37]. Lacking experimental data for whether and at what rate MukBEF complexes might traverse each other, we settled on running simulations with a short stalling time of *τ*_stall_ = 7 seconds (results in the main text), or without any traversal at all, corresponding to the limit *τ*_stall_ ≫ *τ*.

The theoretical work by Goloborodko *et al*. focused on the limit of loop-extruders incapable of traversing each other [44]. In this limit of *τ*_stall_ ≫ *τ*, for λ*/d <* 1, the mean loop length *ℓ* is expected to fall below λ, since loop-extruders stop upon colliding with each other. In the opposite limit, when traversal rate is much faster than the step rate, *τ*_*stall*_ ≪ *d*_step_*/v*_LE_, where *d*_LE_ is the loop-extruder step size, we approach the limit of non-interacting loop-extruders, and the mean loop length *ℓ* is always equal to λ, independent of the loop-extruder density.

In our simulations, we suppose that the loop-extruders have a stepping rate of 0.75 kb/s, similar to *B. subtilis* [10]. On a lattice with *d*_LE_ = 1 kb steps, this gives a stepping time of 1.3 seconds, lower than the traversal time. We hence expect the system to have a sufficiently slow traversal rate that loop-extruder collisions noticeably slow down loop extrusion, and consequently *ℓ <* λ (Supp. Fig. S5). However, traversal events still happen quite frequently.

When bypassing is disabled in our simulations, the mean backbone length increases and the mean loop size decreases (Supp. Fig. S12). This is as expected, since increasing the time that loop-extruders stall upon encountering each other decreases the size of their loops, and after a loop-extruder from a collided pair falls off, the length of the backbone increases more. Since both backbone compaction and a larger loop size stabilise chromosome order, for the same *M* and λ, simulations without loop-extruder bypassing show less stable left-*ori*-right organization than simulations with bypassing (Supp. Fig. S12). However, if the loop-extruder processivity is increased to achieve a similar level of backbone compaction as before, the system can be stabilised. Without bypassing, we find stable left-*ori*-right order for *M* = 50 and λ ≈ 300 kb, or *M* = 30 and λ = 330 kb.

## Appendix C: Simulation time units

Our simulations include two time-scales; the time-scale of the 1D lattice simulations, as well as the time-scale of polymer relaxation in the 3D simulations. Although the time-scale in 1D simulations can be set accurately given experimental data for loop-extrusion rates, the time-scale of polymer simulations must be calibrated against experimental data.

Since we do not know the speed of loop extrusion of MukBEF, we initially assume a loop extrusion speed of 46 kb min^−1^ [10] and a stalling time of *τ*_stall_ = 7 s upon loop extrusion collision [43]. The loop-extruder configurations are saved at time-intervals of *τ*_1D_ = 1 s. We note that if the loop-extruder speed is slower, *τ*_1D_ could simply be interpreted as a larger time-interval. For instance, a loop-extrusion speed of 16-19 kb/min, as seen in *Caulobacter crescentus* [11], would correspond to *τ*_1D_ between 2.4 and 2.9 s.

Each 1D simulation is first allowed to equilibrate for 3000 *τ*_1D_. The simulations are then run and sampled for 120000 *τ*_1D_.

For each 1D simulation, we run a corresponding 3D simulation. After loading a new loop-extruder configuration, the polymer is allowed to relax for 2500 3D simulation time steps. The next loop-extruder configuration is then loaded. We save the polymer configuration at *τ*_3D_ = 60 *τ*_1D_ intervals; this defines the simulation time unit.

To calibrate our polymer simulation time, we track the mean squared displacements (MSD) of loci for 500 *τ*_1D_ in WT simulations at a sampling rate of 1/*τ*_1D_, and compare the results to data on the diffusivity of loci by [78]. Weber et al. found that *E. coli* loci move subdiffusively, such that MSD ≈ 4*D*_app_*t*^0.4^. Our simulations show sub-diffusive motion of loci with a similar exponent (Supp. Fig. S11).

We find that interpreting *τ*_1D_ as 1 − 3 s is consistent with the MSD data by Weber *et al*. (Supp. Fig. S11). This means that the 3D simulations are sampled at intervals equivalent to approximately *τ*_3D_ ≈ 1 − 3 minutes. We note that this is an approximation; parameters such as *M* and λ can affect the simulation MSD curves, and experimentally measured MSD magnitudes are affected by factors such as global transcription and translation rates [78] as well as genomic location [79].

## Appendix D: Defining the orientation angle *θ*

To define the orientation angle *θ* of a given chromosome configuration, we use the Julia package LsqFit [80] to fit the function 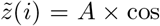 (*i ×* 2*π/N* + *θ* − *π/*2) to the long-axis positions *z*(*i*) of all monomers *i*. To quantify the quality of the fits, we calculated the sum of squared residuals (SSR) values. For statistics of the orientation angle, only fits with an SSR value below 500 are used. Supp. Fig. S2 shows examples of fits with SSR below and above this value.

Even a single flipping event can cause the standard deviation of *θ* over simulations to increase significantly (Supp. Fig. S3). To parameterize the stability of left-*ori*-right order with a measure that is less sensitive to flipping, we therefore consider the orientation angle modulo *π*, such that left-*ori*-right and right-*ori*-left order are both assigned an angle of *θ* = 0. We find that the distribution of *θ* mod *π* shows a relatively flat distribution for simulations with notable diffusion of *θ*, whereas simulations with occasional or no flips show a clear peak at *θ* = 0 (Supp. Fig. S3). We hence use the standard deviation of *θ* mod *π* as a simple measure of left-*ori*-right stability in our simulations.

## Appendix E: Simulations in an infinite tube

To test our predicted scaling for the effective confinement diameter (Eq. (4)), we simulate the system in an infinitely long tube of diameter *D*. We then measure the maximal extension of the looped region along the tube, *z*_bb_. If the extruded loops would not affect the extension of the backbone, we would expect that the extension of the looped region would scale linearly with the backbone length; *z*_bb_ *∝ N*_bb_. On the other hand, if the loops affect the backbone’s effective confinement diameter, we would expect 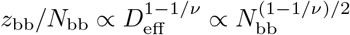.

Plotting *z*_bb_*/N*_bb_ against *N*_bb_ shows an approximate scaling law. However, simulations with 100 loop-extruders seem to show a higher than expected extension (Supp. Fig. S5). We note that *N*_bb_ does not reflect the true contour length of the backbone between the first and last loop in the looped region, and we therefore define a more accurate measure.

To calculate the effective contour length of the backbone of the looped region, we start by calculating the mean number of monomers between loops in the looped region, *Ñ* _bb_. We distinguish this from *N*_bb_, since we only count backbone monomers that fall between the first and last loop. The excluded monomers can be considered an extension of the unlooped region.

Next, we note that each loop-extruder at the base of a loop contributes to the backbone length by the SMC size *ℓ*_SMC_. We therefore calculate the mean number of baseloops (excluding loops within loops), *M*_base_, and define the effective contour length of the backbone as

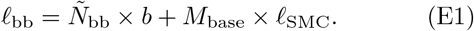

In practice, including the contribution from the SMC length becomes more relevant as the number of loop-extruders increases. Using this effective contour length, we find that 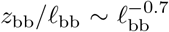 (Supp. Fig. S5). This exponent is consistent with *ν* ≈ 0.42, lower than expected for a linear polymer, and closer to the value 2*/*5 expected for ring polymers. Some differences from scaling theory might be expected both due to finite size effects as well as the finite excluded volume potential used in the simulations. Nevertheless, the results confirm that the expected extension of the looped region mainly depends on the backbone contour length.

## Appendix F: Mushroom regime

To estimate whether or not our system falls into the mushroom regime, we need to approximate both the loop spacing and the loop radius of gyration.

A simple approximation for the genomic loop spacing is given by *N*_bb_*/M*. In reality, this gives an underestimate, since multiple loop-extruders can be nested in the same loop. Using Eq. (1), for *M* = 50 and λ = 30, 150, 300, we find *N*_bb_*/M* ≈ 50, 10, 1.5 correspondingly. Using *b* = 30 nm, these values give maximal extended loop spacings of 300, 150 or 45 nm.

For ring polymers in a melt, Cates & Deutsch used a Flory argument to predict that the rings’ radius of gyration should scale with an exponent of approximately *ν* = 2*/*5, lower than for linear polymers [55]. More recent work suggests that this exponent corresponds to a transition regime, and that at higher loop densities *ν* ≈ 1*/*3 [60, 61]. Using the larger value *ν* = 2*/*5 as a conservative estimate, we find *R*_g_ = 120, 220, 290 nm for λ = 30, 150, 300 correspondingly. Hence our calculation suggests that for all but the highest processivities 300 kb, the loop spacing is of the same order of magnitude as the loops’ radius of gyration, and the chromosome is expected to be in the mushroom regime. This implies that the loop extension should scale as 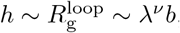, as seen in our simulations (Supp. Fig. S5).

Appendix G: Maximum number of loop-extruders that can maintain stability

To explain why 200 loop-extruders were unable to maintain left-*ori*-right order in our simulations, we derive an upper limit for the number of loop-extruders that can give rise to sufficiently large loops such that *h > D/*4.

Suppose that the system is still in the mushroom regime, such that *h* ∼ *R*_g_ ∼ *bℓ*^*ν*^. We hence find a minimum loop size *ℓ*_min_ ∼ (*D/*4*/b*)^1*/ν*^ required for stable left-*ori*-right order. For *D* = 1 µm, *b* = 0.03μm and *ν* = 2*/*5, this gives *ℓ*_min_ ∼ 200 kb. This is in line with our simulations, where stability for *M* = 30, 50, 100 requires processivities between 125-300 kb.

In the sparse and intermediate density regime, with λ*/d* ≲ 1, the minimum loop size can be reached provided that λ ≈ *ℓ*_min_. However, at high loop-extruder densities, when λ*/d* ≳ 10, the mean loop length scales as *ℓ* ∼ *d* log(λ*/d*) ≪ λ [44], and the minimum loop size becomes impossible to reach for reasonable λ. The condition for instability hence becomes *ℓ*_min_ ∼ 10 *d*; or equivalently, the number of loop-extruders should reach 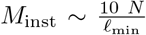. Using *N* = 3800 kb and *ℓ*_min_ ∼ 200 kb, this gives *M*_inst_ ∼ 190. Although the scaling argument neglects prefactors of order unity, the result agrees with our simulations that show stable left-*ori*-right organization for *M* = 100 but not *M* = 200.

## Appendix H: Simulations with fixed loop-extruders

To confirm that activity of loop-extruders is not essential to the maintenance of left-*ori*-right order in our model, we conduct simulations where we sample a single loop-extruder configuration from the 1D simulations, and then perform a 3D simulation with loop-extruders at these fixed positions. Our results show that fixed loop-extruders are capable of maintaining left-*ori*-right order, as seen in simulations with active loop extrusion (Supp. Fig. S10). This confirms that the main role of loop extrusion in our model is to give rise to a dense set of large loops on the chromosome; loop-extruder activity is not necessary for stable left-*ori*-right order.

## Notes

### Competing Interest Statement

The authors have declared no competing interest.

