## Supplementary Figures for "Loop-extruder mediated rigidity can globally order bacterial chromosomes"

Janni Harju[1], Till Armbruster[1,2], and Chase P. Broedersz[1]

[1]Department of Physics and Astronomy, Vrije Universiteit Amsterdam, 1081 HV Amsterdam, The Netherlands
[2]Institute for Theoretical Physics, Utrecht University, Princetonplein 5, 3584 CC Utrecht, The Netherlands

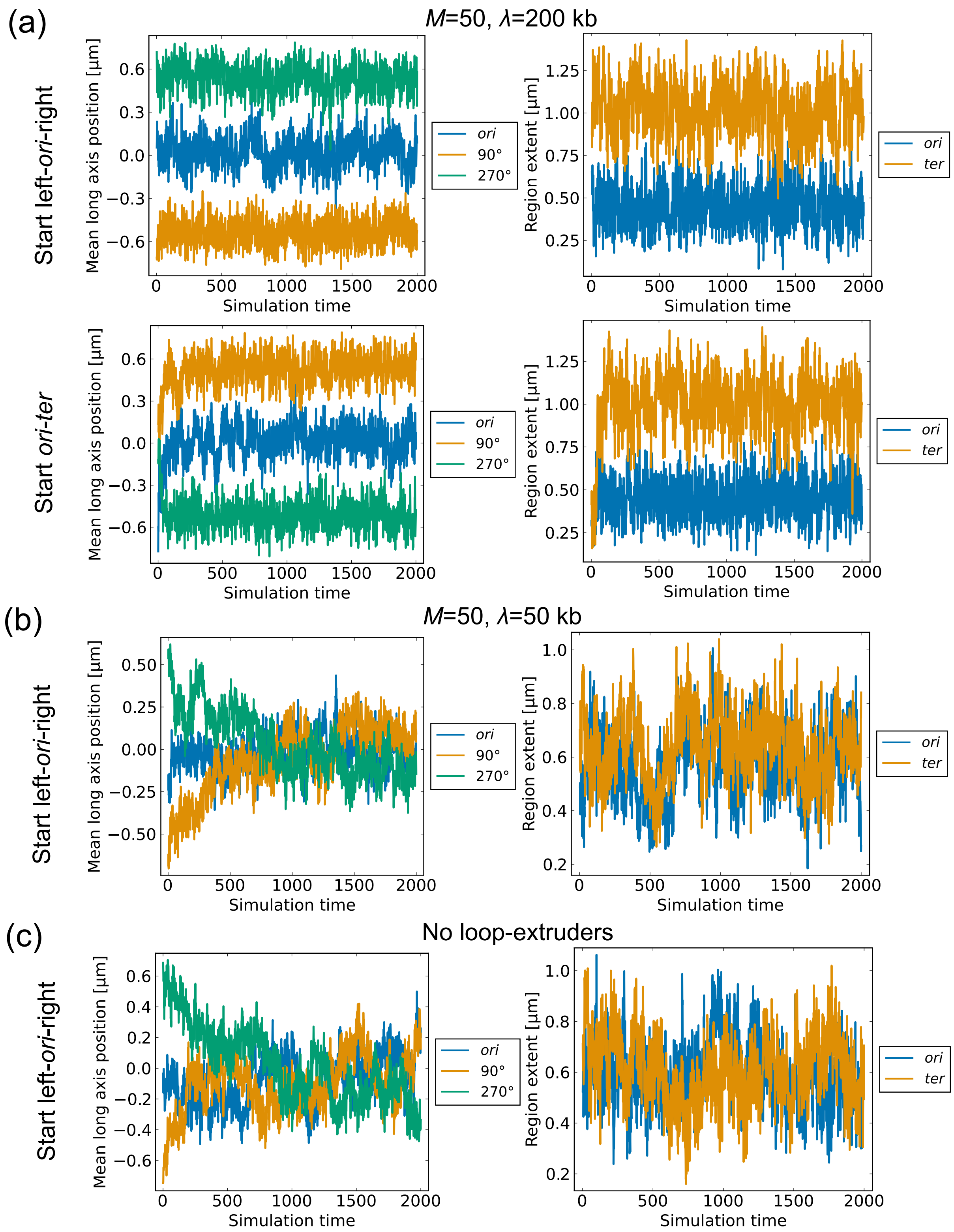


Supplementary Figure S1: **Simulation convergence.** **(a)** Table showing convergence of mean long axis positions and mean long axis extensions of 800 kb regions as a function of simulation time. First row: simulations starting with a left-*ori*-right configuration. Second row: simulations starting with an *ori*-*ter* configuration. In the second case, we flip configurations so that the left arm has negative long axis positions. Otherwise, due to the random orientation of the chromosome, the mean positions would converge to zero. **(b)** Similar to (a), but for simulations with $\lambda = 50$ kb. The loop size is insufficient for left-*ori*-right order. **(c)** Similar to (b), but for simulations without loop-extruders.

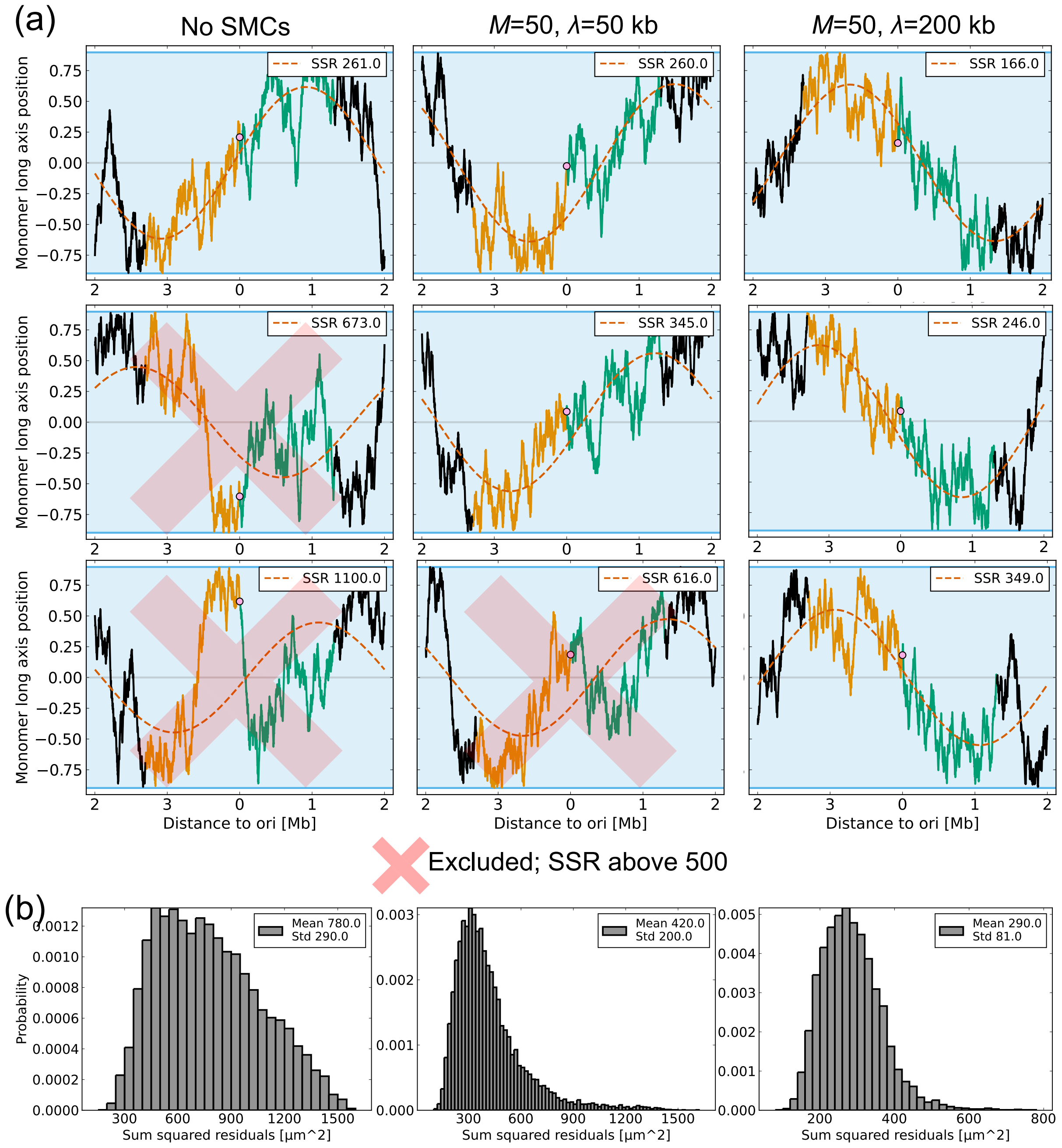


Supplementary Figure S2: **Examples of fitted sinusoidal curves. (a)** Table of examples of sinusoidal fits to the long axis positions of loci. Red crosses indicate fits with a sum of squared residuals (SSR) value above 500 μm$^2$, which were excluded from the angle trajectories. **(b)** Histograms of the SSR values for simulations without loop-extruders, and with $M = 50$, $\lambda = 50$ or 200 kb.

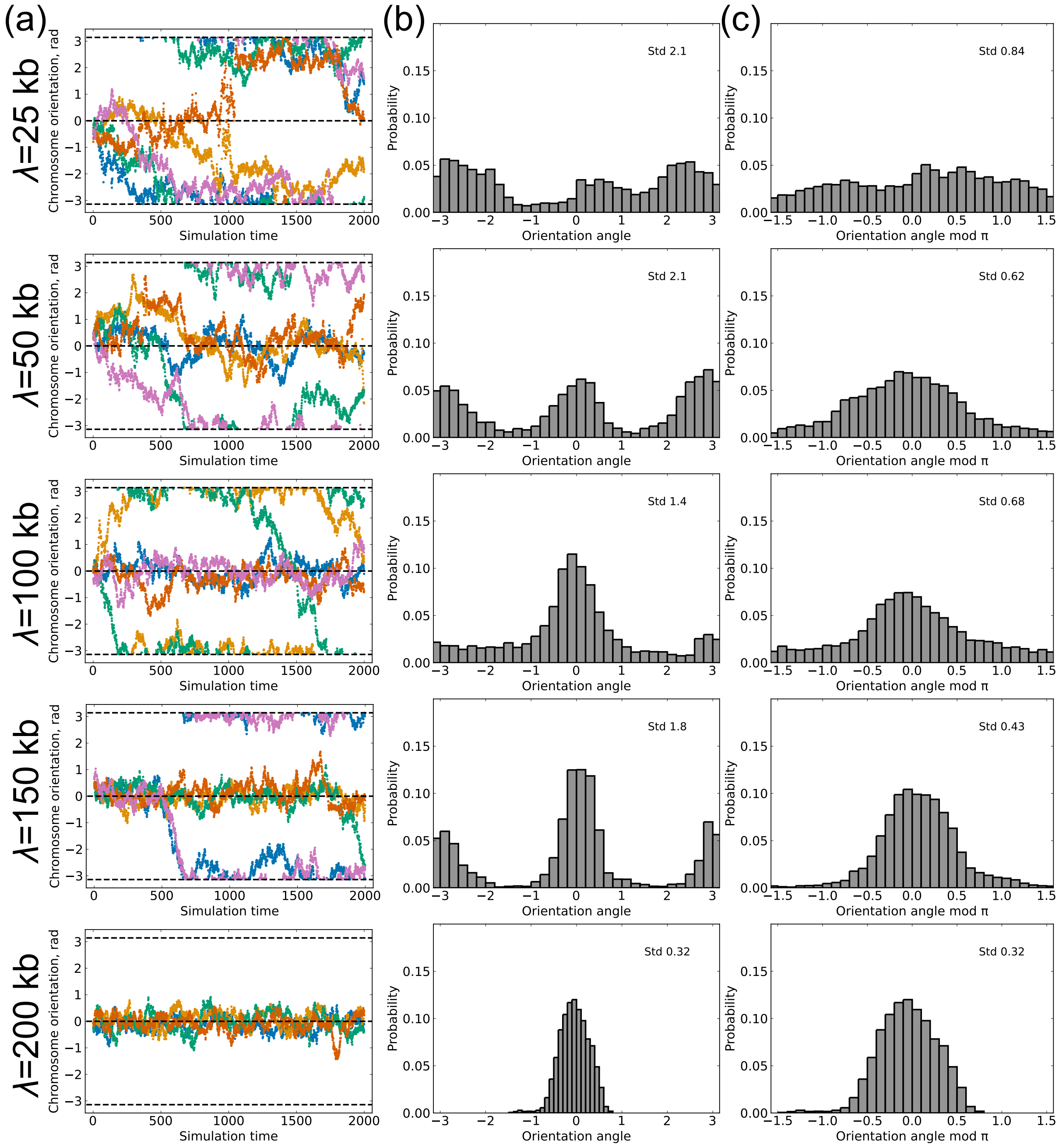


Supplementary Figure S3: **Table of $\theta$ distributions for varying processivities.** **(a)** Trajectories of the orientation angle over time. $\lambda$ for each row indicated on the left, $M = 50$ for all shown plots. **(b)** Histograms of the orientation angle. For simulations with occasional flipping, two peaks corresponding to left-*ori*-right or right-*ori*-left order can be seen. **(c)** Histograms of the orientation angle modulo $\pi$.

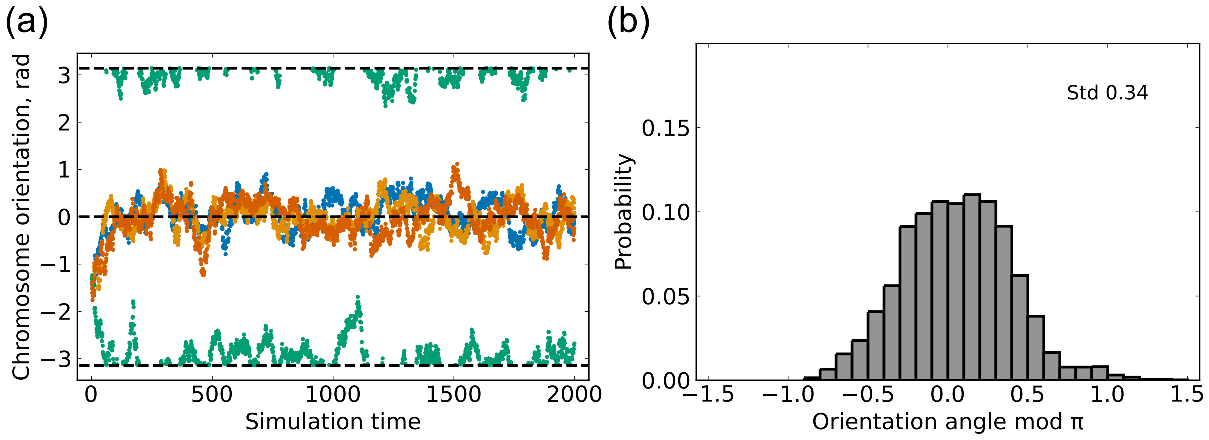


Supplementary Figure S4: **Simulations starting from *ori*-*ter* configuration.** **(a)** Orientation angle trajectories for simulations starting from an *ori*-*ter* configuration, with $\theta = -\pi/2$. The angles converge to either left-*ori*-right or right-*ori*-left in less than 200 simulation time steps. Used $M = 50$, $\lambda = 299$ kb. **(b)** For time-points $> 1000$, the distribution of orientation angles modulo $\pi$. Standard deviation comparable to simulations starting from a left-*ori*-right orientation (Supp. Fig. S3 (c)).

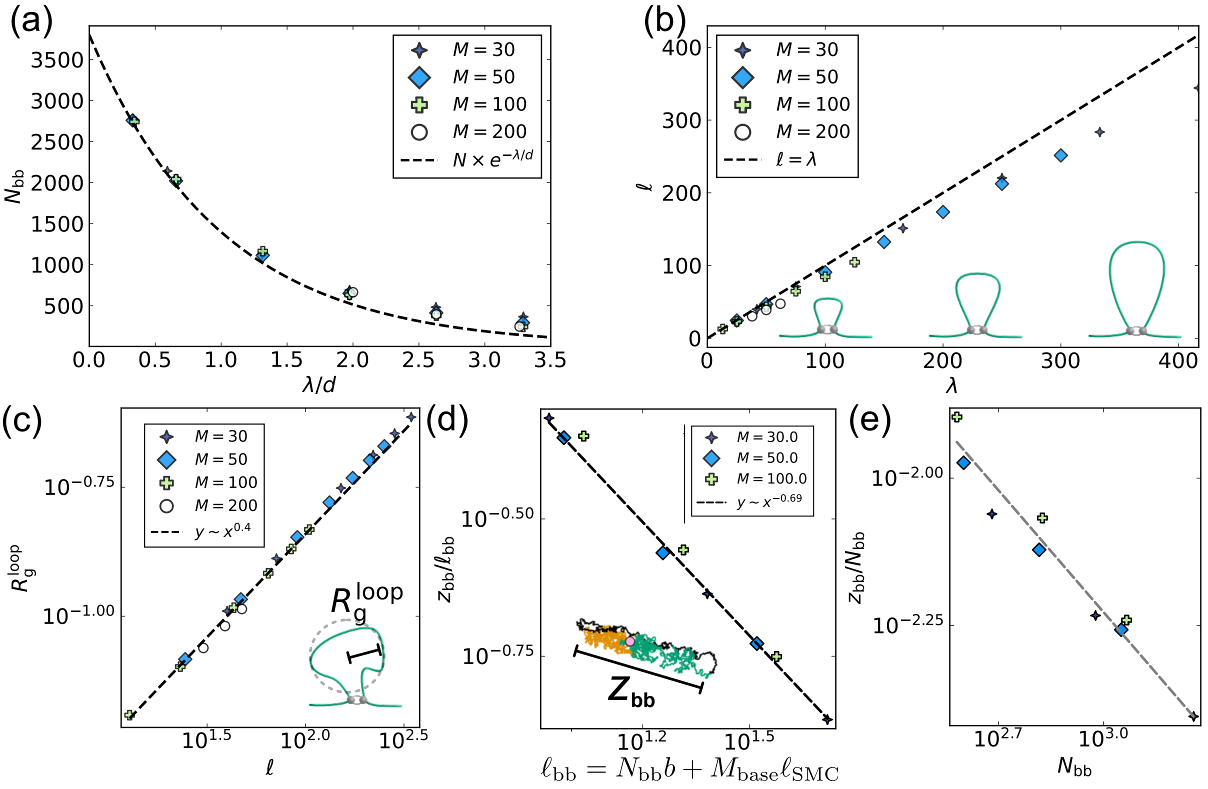


Supplementary Figure S5: **Confirming scaling regimes in simulations.** **(a)** The scaling of the backbone length $N_{\mathrm{bb}}$ with the processivity divided by the mean loop-extruder spacing. Prediction shows theoretical estimate based on work by Goloborodko *et al.* (1). **(b)** The scaling of the mean loop length $\ell$ with the processivity $\lambda$. Collisions between loop-extruders result in $\ell < \lambda$. **(c)** The scaling of the loop radius of gyration with the mean loop length. Line indicates scaling of $R_{\mathrm{g}}^{\mathrm{loop}} \sim \ell^{\nu}$, with $\nu = 2/5$ as often seen for solutions of ring polymers. **(d)** Scaling of the extension of the looped region with the effective genomic length of the backbone in simulations in an infinite tube. The simulations show an approximate scaling consistent with $z_{\mathrm{bb}}/\ell_{\mathrm{bb}} \sim D_{eff}^{1-1/\nu} \sim \ell_{\mathrm{bb}}^{(1-1/\nu)/2}$, with $\nu \approx 0.42$. **(e)** Similar to (d), but using the mean number of monomers in the backbone, $N_{\mathrm{bb}}$. More deviations from the approximate scaling are visible. Exponent is the same 0.69 as in subfigure (d).

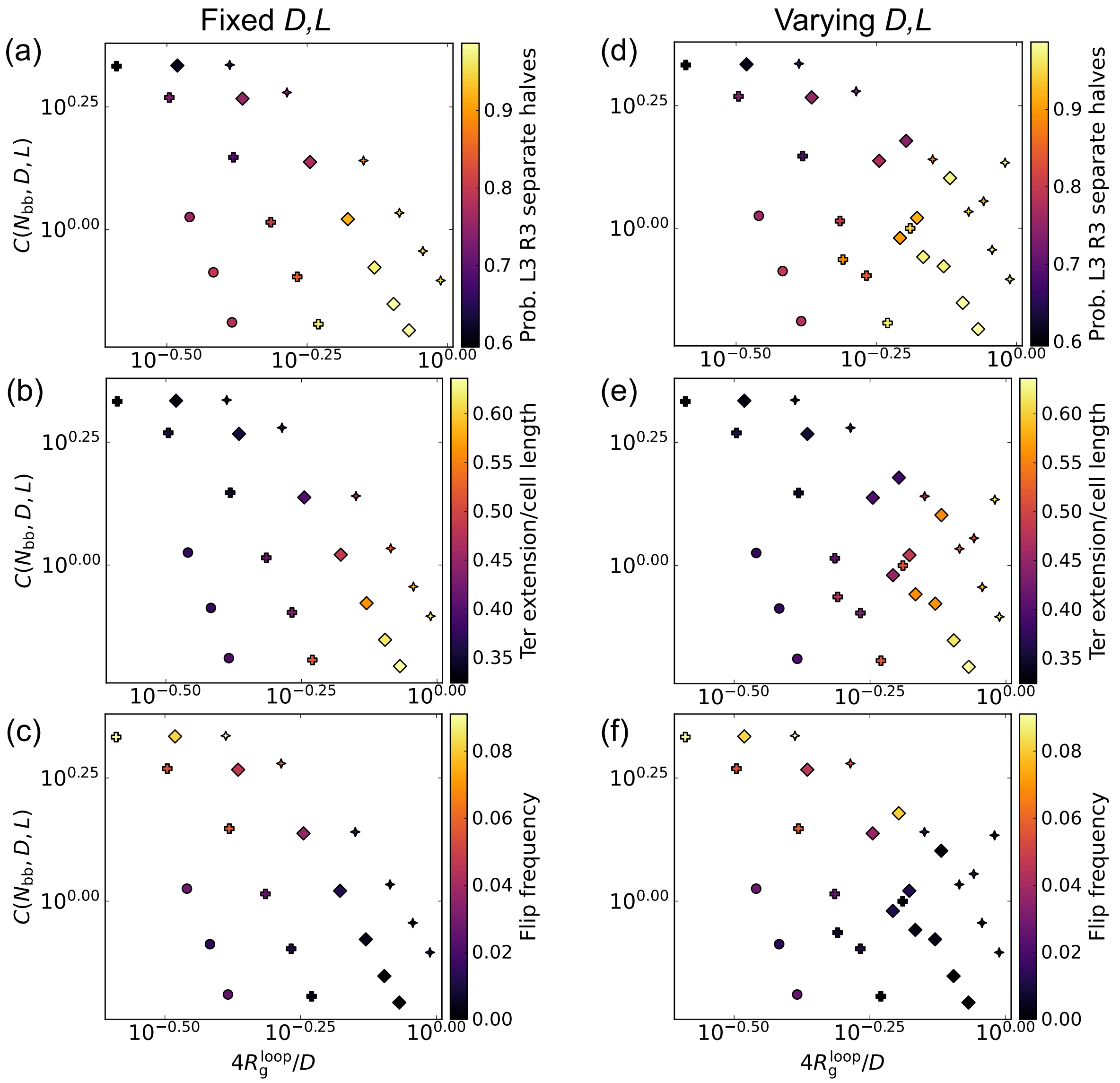


Supplementary Figure S6: **Phase diagrams using other measures of left-*ori*-right order.** **(a)** Similar to Main Figure 3(g), but using the probability of finding the L3 and R3 loci in different nucleoid halves. **(b)** Using the end-to-end distance of the unlooped *ter* region relative to the confinement length. **(c)** Using the frequency at which the L3 and R3 loci switch their relative long axis positions. **(d-f)** Similar to (a-c), but including data from simulations with different confinement dimensions.

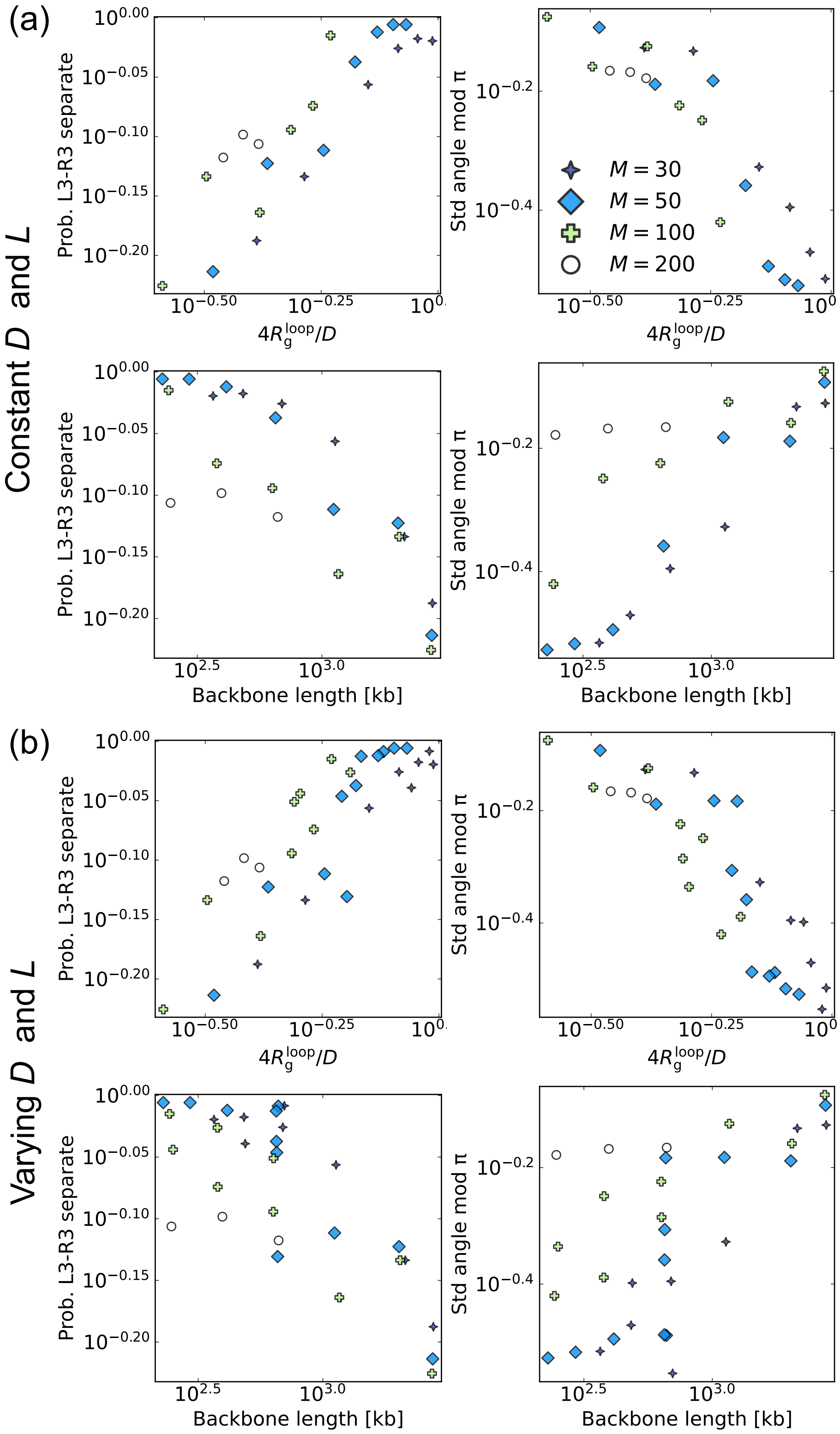


Supplementary Figure S7: **Single parameters do not explain organizational stability.** **(a)** Scatter plots of organizational stability measures against either the loop size relative to the confinement diameter (top) or the length of the backbone (bottom) from simulations with varying $M$ and $\lambda$. The curves do not collapse, since stability requires both compaction and a sufficient loop size. **(b)** Similar to **(a)**, but including simulations with varying confinement dimensions $D$ and $L$.

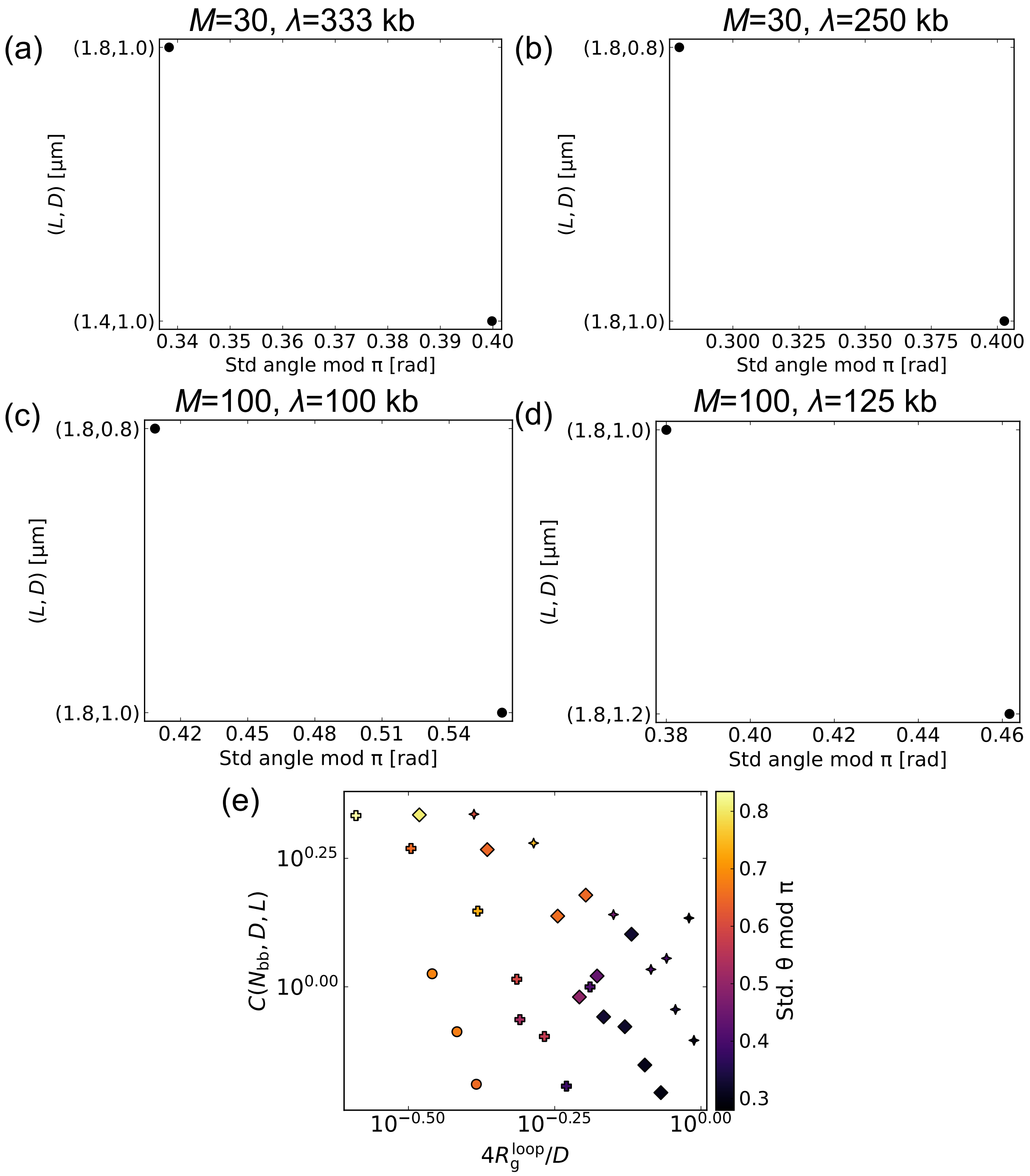


Supplementary Figure S8: **Longer and narrower confinement increases left-*ori*-right stability.** Data shown for simulations with varying $M$ and $\lambda$, as indicated in the figure. **(a)** Shortening the confinement decreases left-*ori*-right order stability. **(b-c)** Narrowing the confinement increases stability. **(d)** Widening the cell decreases stability. **(f)** Similar to Main Figure 3 (g), but including results from simulations with varying confinement dimensions.

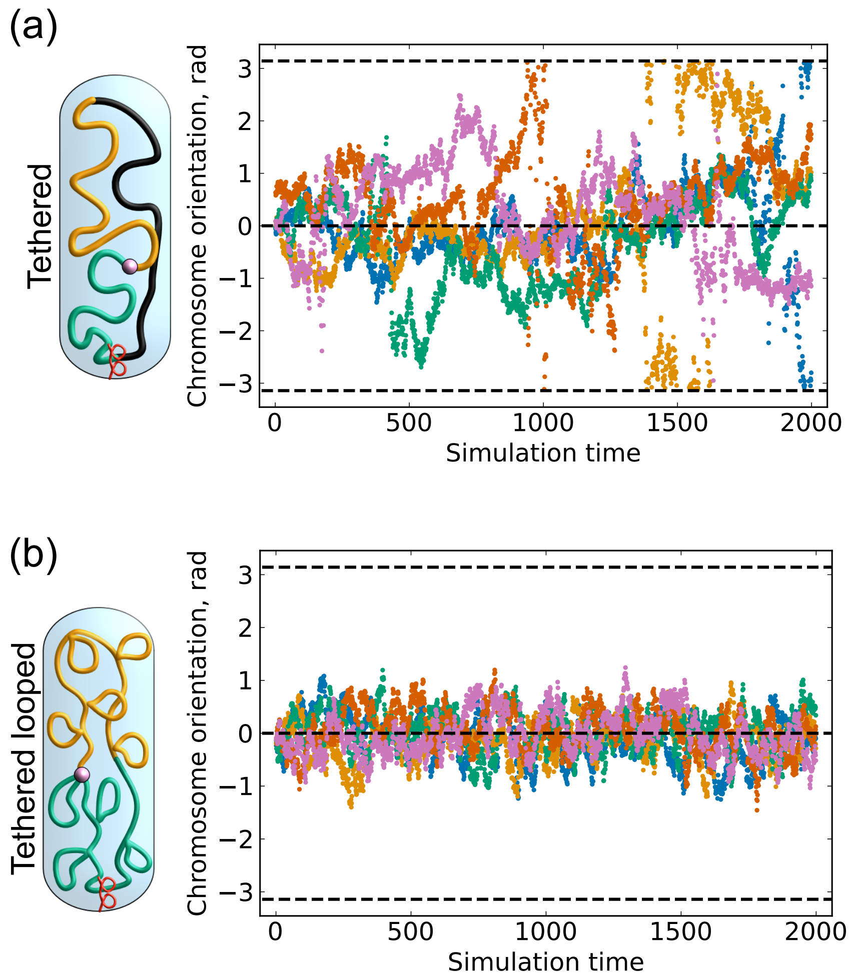


Supplementary Figure S9: **Effects of anchoring a locus to the nucleoid pole. (a)** Fitted orientation angle trajectories for tethered model without loop-extruders. Breaks in the angle trajectories indicate time-points when the long axis position curve was not well-fitted with a sinusoidal function (See Supp. Fig. S2 for details). **(b)** Similar to (a), but for the tethered model with $M = 50$ loop-extruders, $\lambda = 200$ kb, but no unlooped *ter* region.

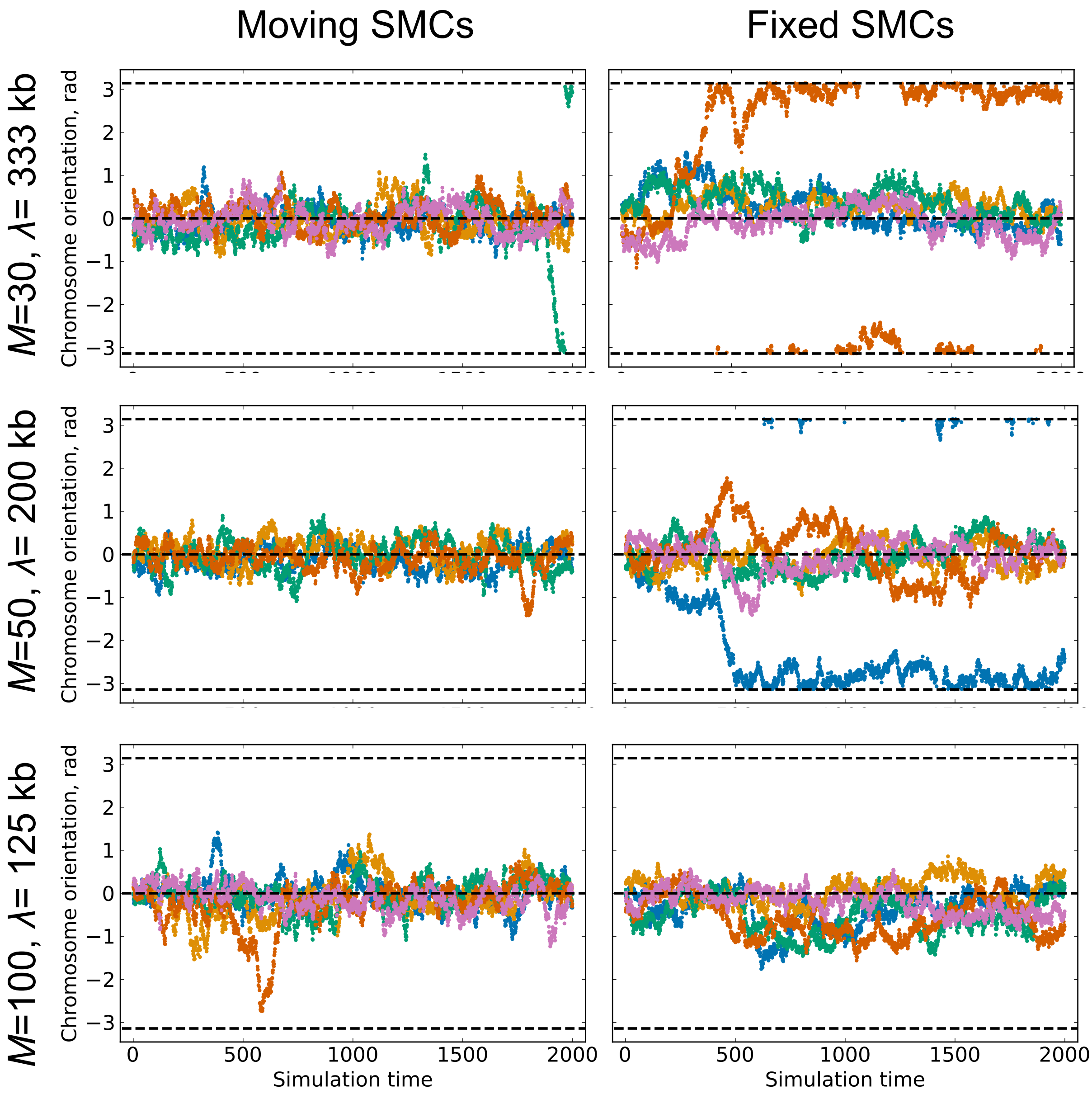


Supplementary Figure S10: **Dynamic loop-extruder is not required for organizational stability.** A table of orientation angle trajectories with either dynamic loop-extrusion, or loop-extruders that were held at fixed positions for the duration of a 3D simulation. Fixing loop-extruders only slightly destabilises left-*ori*-right orientation.

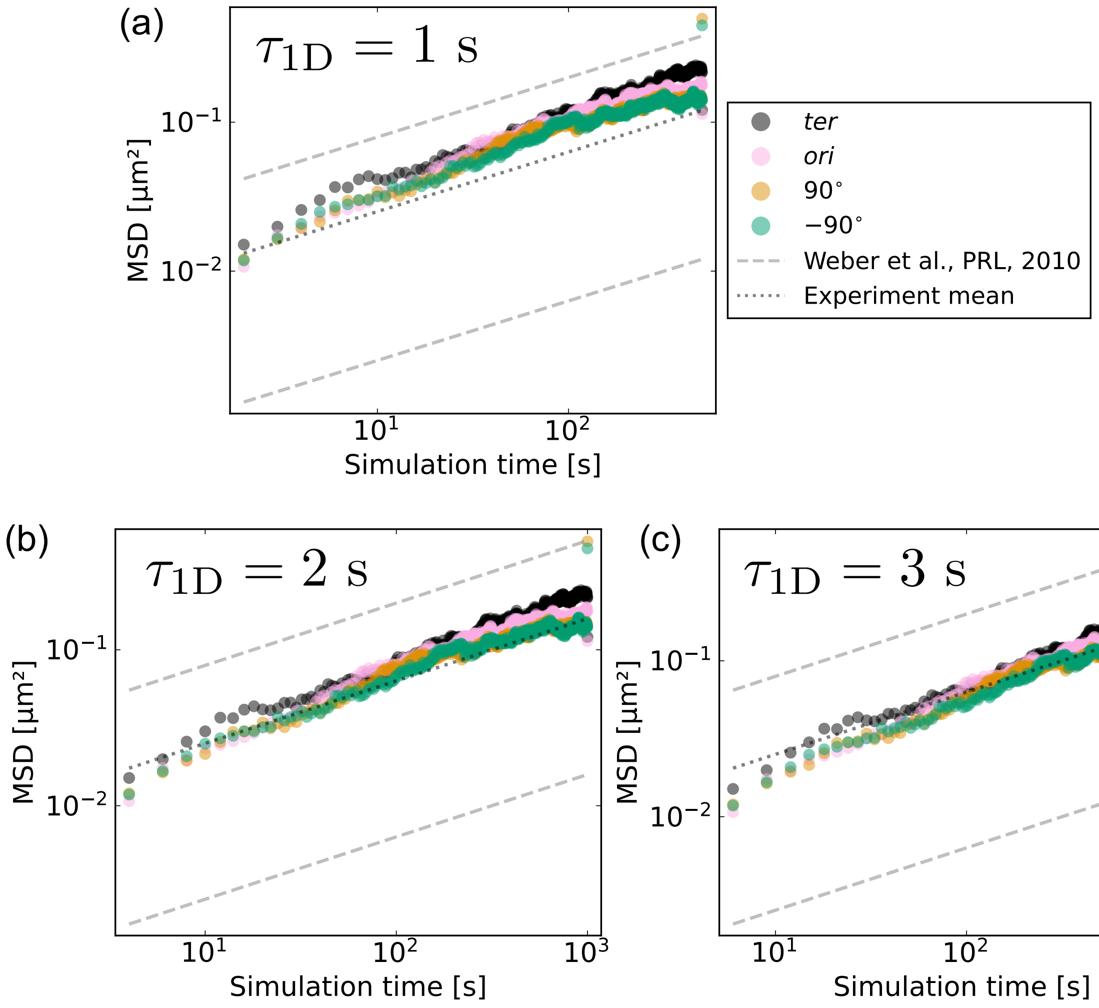


Supplementary Figure S11: **Interpreting simulation time.** **(a)** The MSD of loci as a function of time, assuming $\tau_{1D} = 1$ s, corresponding to a loop-extrusion speed of 46 kb/s, and a 3D simulation time unit of 1 min. Dashed lines indicate the scaling of MSD$\propto t^{0.4}$ observed by Weber *et al.* (2), as well as the range of and mean coefficient seen in experiments. The simulation data shows slightly faster dynamics than expected. **(b)** Similar to (a), but assuming $\tau_{1D} = 2$ s, corresponding to a loop-extrusion speed of 23 kb/s, and a 3D simulation unit of 2 min. At short times, the dynamics now appears to be close to the experiment mean. **(c)** Similar to (a), but assuming $\tau_{1D} = 3$ s, corresponding to a loop-extrusion speed of 15 kb/s, and a 3D simulation unit of 3 min. At short times, the dynamics now appears to be somewhat too slow.

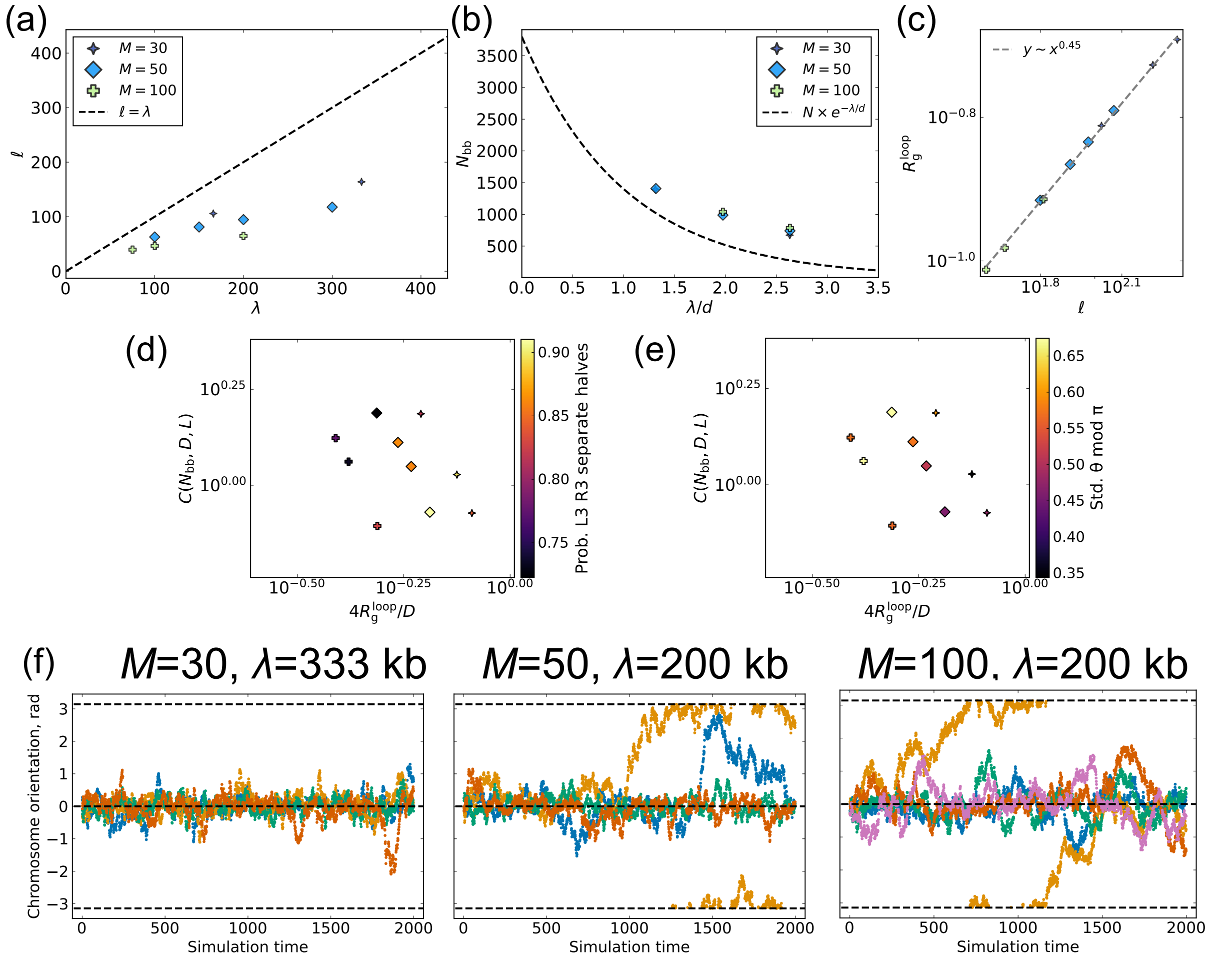


Supplementary Figure S12: **Simulations without loop-extruder bypassing.** **(a)** Mean loop-size decreases when loop-extruders cannot bypass each other after colliding. **(b)** The mean backbone length increases without bypassing. **(c)** Without bypassing, the loop radius of gyration scales with a fitted exponent of $\nu \approx 0.46$. **(d)** Probability of L3 and R3 loci being in separate nucleoid halves, as a function of the loop size and compaction factor. For the same processivities, since without bypassing both the loop size decreases and the backbone length increases, simulated systems are in a less stable regime than with bypassing. **(e)** Similar to (d), but with the standard deviation of the orientation angle modulo $\pi$. **(f)** Example orientation angle trajectories from simulations without loop-extruder bypassing.
